# Population genomic consequences of novel life history and mating system adaptation to a geothermal soil mosaic in yellow monkeyflowers (*Mimulus guttatus*)

**DOI:** 10.1101/2021.07.29.453448

**Authors:** Kory M. Kolis, Colette S. Berg, Thomas C. Nelson, Lila Fishman

## Abstract

Local selection can promote phenotypic divergence despite gene flow across habitat mosaics, but adaptation itself may generate substantial barriers to genetic exchange. In plants, life-history, phenology, and mating system divergence have been particularly proposed to promote genetic differentiation in sympatry. In this study, we investigate phenotypic and genetic variation in *Mimulus guttatus* (yellow monkeyflowers) across a geothermal soil mosaic in Yellowstone National Park (YNP). Plants from thermal annual and nonthermal perennial habitats were heritably differentiated for life history and mating system traits, consistent with local adaptation to the ephemeral thermal-soil growing season. However, genome-wide genetic variation primarily clustered plants by geographic region, with little variation sorting by habitat. The one exception was an extreme thermal population also isolated by a 200m geographical gap. Individual inbreeding coefficients (FIS) were higher (and predicted by trait variation) in annual plants and annual pairs showed greater isolation by distance at local (<1km) scales. Finally, YNP adaptation does not re-use a widespread inversion polymorphism diagnostic of annual vs. perennial *M. guttatus* range-wide, suggesting a novel genetic mechanism. Overall, this work suggests that life history and mating system adaptation strong enough to shape individual mating patterns does not necessarily generate incipient speciation without geographical barriers.

## Introduction

The opposition between gene flow and selection across heterogeneous landscapes governs both local adaptation and the emergence of new species. Extrinsic barriers to gene flow (e.g., distance, dispersal barriers, or gaps in suitable habitat) allow allopatric populations to drift or adapt independently to speciation (reviewed in Gavrilets 2003; Coyne and Orr 2004). However, a small number of migrants between populations can be enough homogenize genetic variation genome-wide (Slatkin 1973; Lenormand 2002), even if environmental gradients or mosaics impose divergent selection. At one extreme, this interplay between selection and gene flow may result in a handful of adaptive outlier loci standing out against a background of low genome-wide differentiation (Jones et al. 2018). At the other, local adaptation itself may generate strong barriers to gene flow among habitats, extending genetic differentiation beyond the loci under divergent selection and potentially initiating speciation. Theory shows that speciation in the absence of geographic barriers is particularly possible when adaptation involves so-called “magic traits”, which simultaneously confer habitat-specific fitness and enforce strongly assortative mating (Smadja and Butlin 2011; Servedio et al. 2011; Kopp et al. 2018). However, while there is overwhelming genomic evidence that adaptation and speciation often progress in contexts including gene flow, it remains ambiguous whether and how local adaptation can generate reproductive isolation and genetic differentiation in the absence of geographic barriers.

In flowering plants, soil mosaics provide an ideal context in which to jointly consider adaptation and speciation without geographic barriers (reviewed in Rajakaruna 2018). Differences in soil drying regimes, nutrient availability, and mineral concentrations between geologically-distinct soils can impose strong divergent selection (Brady et al. 2005), often over distances within the range of seed and pollen movements (Richardson et al. 2014). Selection across such edaphic mosaics may result in differentiation of key adaptive traits controlled by major loci, but little neutral genomic divergence, as the classic blue-white flower polymorphism in *Linanthus parryae* (Schemske and Bierzychudek 2001; 2007) or copper tolerance in the yellow monkeyflower *Mimulus guttatus* (Lee and Coop 2017). Nonetheless, numerous edaphically-specialized endemic plant species suggest that soil adaptation may often contribute to speciation over short spatial scales, potentially by driving divergence in pollination, mating system, or phenological traits (Rajakaruna 2018, Baack et al. 2015). For example, shifts in flowering time associated with adaptation to a volcanic/nonvolcanic soil mosaic have been invoked as the key to sympatric speciation in island palms (Savolainen et al. 2006). The costs of maladaptive gene flow across sharp habitat boundaries may even directly favor shifts in reproductive biology that reinforce isolation, as has been posited for sweet vernal grass adapting to toxic mine tailings (Antonovics 2006) and long-term fertilization experiments (Silvertown et al. 2005). Teasing apart the factors that result in these different outcomes requires a complete picture of how natural selection, ecological distance, and geography contribute to genome-wide patterns of differentiation during edaphic adaptation.

We investigate how life-history and mating system divergence influence population structure and individual genomic variation in yellow monkeyflowers across a geothermal soil mosaic in Yellowstone National Park (YNP). The yellow monkeyflowers of the *Mimulus guttatus* species complex are a model system for understanding the evolutionary genetics of edaphic adaptation (MacNair 1983; Ferris et al. 2014; Lee and Coop 2017; Selby and Willis 2018), as well as life history (Lowry and Willis 2010; Oneal et al. 2014; Twyford and Friedman 2015), phenological evolution (Friedman and Willis 2013; Fishman et al. 2014; Kooyers et al. 2015), and mating system divergence (Fishman et al. 2002), creating a robust comparative framework. Across a latitudinal range from Alaska to Baja California, *M. guttatus* populations exhibit alternative annual and perennial ecotypes, with annuals found in ephemeral wet soils and perennials wherever soils remain saturated through the summer (Lowry and Willis 2010; Oneal et al. 2014; Twyford and Friedman 2015). Numerous traits associated with life history (including flowering time, flower and plant size, and stolon production) map to a diagnostic chromosomal inversion on Chromosome 8 of the *M. guttatus* reference genome (*DIV1*; Hall et al. 2006; Lowry and Willis 2010), and widespread annuals and perennials exhibits strongly elevated differentiation across this large region of suppressed recombination (Oneal et al. 2014; Twyford and Friedman 2015; Gould et al. 2017). Within annual *Mimulus guttatus*, several distinct species have arisen via the evolution of obligate selfing (tiny corollas and no stigma-anther separation, with self-pollination prior to corolla opening) (Fenster 1994; Brandvain et al. 2014). The most widespread, *M. nasutus*, occurs in sympatry with *M. guttatus* across a broad range, occupying ephemerally wet micro-sites that favor spring-flowering (Martin and Willis 2007; Kenney and Sweigart 2016); others (e.g., *M. laciniatus*, *M. cupriphilus*) are endemic to distinct edaphic substrates (MacNair and Cumbes 1989; Ferris et al. 2014). Edaphic adaptation (e.g., to toxic serpentine and copper mine soils) has also occurred within *M. guttatus* without mating system divergence and/or speciation (MacNair 1983; Lee and Coop 2017; Selby and Willis 2018). Thus, edaphic, life history and mating system adaptation have distinct outcomes even within this one species complex.

In Yellowstone National Park (home to >50% of Earth’s geysers), the geothermal influence recapitulates on a microgeographic scale the soil conditions that shape adaptation across the range of *M. guttatus*. Yellow monkeyflowers are one of a few vascular plants that live in extreme geothermal crusts, which create a unique ground-level oasis of winter-annual habitat when cooled by melting winter/spring snow but become deadly hot (45-60°C) and dry by early summer (Lekberg et al. 2012). Throughout YNP, summer-flowering *M. guttatus* also occur in nonthermal wetland habitats (bogs, riversides) that are under snow much of the year, and in diverse intermediate habitats created by geothermally influenced water sources (Fig. 1A). Previous work in this system has focused on two sites, Agrostis Headquarters (AHQ) and Rabbit Creek (RBC), which encompass similar thermal and nonthermal extremes but differ in their degree of habitat-associated population genetic structure (Lekberg et al. 2012; Hendrick et al. 2016). In common gardens, plants from focal thermal sites exhibited shifts in floral morphology and growth form consistent with adaptation to the narrow spatial and temporal window for growth and reproduction in this extreme habitat (but life history traits were not measured; Lekberg et al. 2012). The potential for microgeographic local adaptation in YNP creates an ideal opportunity to understand how traits, environment, and geography together shape genetic variation across complex habitat mosaics.

**Fig. 1.**
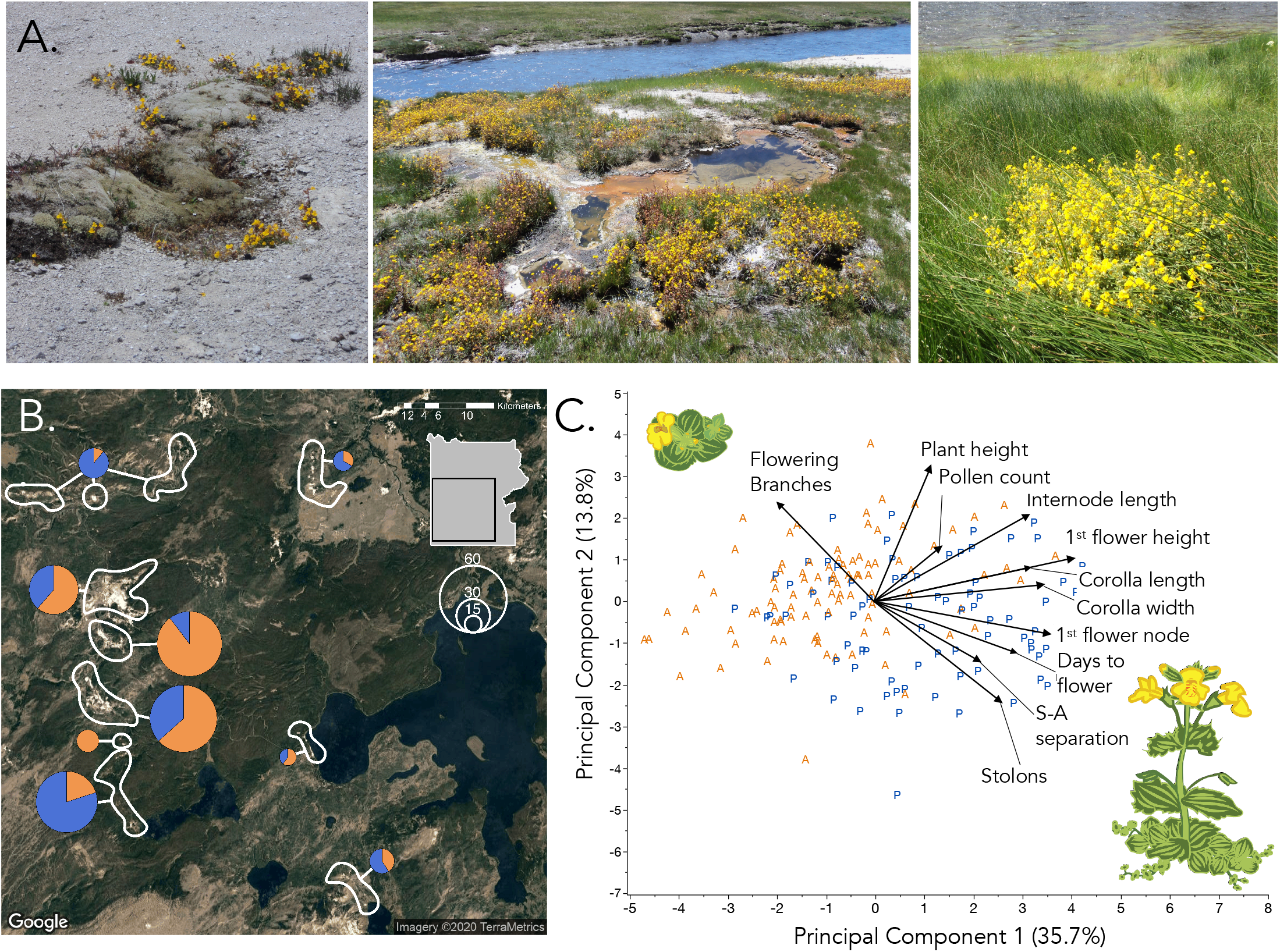
*Mimulus guttatus* exhibits heritable phenotypic variation associated with a geothermal soil habitat mosaic in Yellowstone National Park. **A)** Example habitats, including (left to right) geothermal crusts where precipitation is the only source of water, the fringes of hot pools and streams, and densely-vegetated nonthermal bogs. **B)** For population genetic and phenotypic analyses, *M. guttatus* (total N = 318) plants were collected from habitats defined as summer-dry annual (orange) and summer-wet perennial (blue) habitats in most of the major geyser basins (pies). **C)** The progeny means of plants from thermal annual (A, orange) and nonthermal perennial (P, blue) habitats were differentiated along major principal components defined by life history (stolons vs. flowering branches) and mating system (stigma-anther separation), and plant phenology, architecture, and size. See Table 1 for habitat means and trait heritabilities. Drawings by Mariah McIntosh.

In this study, we characterize heritable phenotypic variation, genome-wide population differentiation, and genetic diversity of plants sampled across the complex matrix of *Mimulus guttatus* habitat in Yellowstone National Park. We defined each collection site as either an annual or perennial habitat (AH and PH, respectively) based on summer dry-down and grew progeny from the wild plants in greenhouse common gardens to extract heritable variation for life history, phenological and floral traits. We used a reduced representation sequencing protocol (ddRADSeq) to genotype the wild individuals at thousands of loci genome-wide, then examined relationships between patterns of variation and habitat, geography, and phenotype. First, we assess whether plants from AH sites are morphologically annual (i.e., do not produce stolons in a common garden), paralleling the heritable differentiation between widespread annual and perennial ecotypes found elsewhere in the species’ range, and characterize the full spectrum of phenotypes from AH and PH habitats. Second, we characterize patterns of genetic variation across the landscape, assign individuals to population clusters, and ask how environment (AH vs. PH) and geographic distance shape diversity within and among clusters. Third, we ask how habitat (and associated phenotypes) affects spatial scale of isolation-by-distance and what heritable phenotypes predict individual homozygosity (FIS, inbreeding coefficient). Finally, although our focus here is not on identifying the genetic basis of adaptive trait divergence, we examine patterns of AH-PH differentiation genome-wide to test whether life history adaptation in Yellowstone thermal-soil habitats re-uses widespread annual-perennial chromosomal polymorphisms or has an independent genetic basis, and then confirm the inference of independence with targeted linkage mapping.

## Materials & Methods

### Study system

*Mimulus guttatus* (Phrymaceae, also known as *Erythranthe guttata*) is an herbaceous plant common in diverse wet soil habitats throughout western North America. Here, we focus on *M. guttatus* variation within Yellowstone National Park (YNP), on the far eastern edge of the species range. In YNP, *M. guttatus* plants are abundant across a complex edaphic mosaic generated by geothermal activity warming soils in winter and making them inhospitable in summer (Fig. 1). Previous work characterized genetic and phenotypic variation at just two focal sites with adjacent extremes (Agrostis Headquarters; AHQ) and Rabbit Creek (RC) (Lekberg et al. 2012; Hendrick et al. 2016). Here, we sampled *M. guttatus* in diverse nonthermal and thermal habitats in all major geyser basins except Mammoth and Mud Volcano, where high and low pH, respectively, may inhibit *M. guttatus* occurrence.

### Field collections and habitat assignment

*M. guttatus* individuals and habitat data were collected from >150 geo-referenced sites across YNP’s geyser basins (n = 2-3 plants ~1m apart per site, Fig.1b, Supplemental Table S1). We visited each site twice, timed as logistically feasible. The first visit (Spring/Summer 2017) was as early in the site-specific growing season as possible to collect adult plants for transplanting into a greenhouse for DNA collection and generation of progeny (see below). We attempted to sample individuals from both potentially annual (e.g., dry, highly thermally influenced) and perennial habitats within each region. In August 2018, we re-visited and categorized each collection location as annual habitat (AH) or perennial habitat (PH) based on soil moisture and the presence of live *M. guttatus*: AH = dry with dead plants; PH = wet with live adult plants.

### Common garden phenotyping

To extract heritable phenotypes from our field plants under common garden conditions, we hand-pollinated one plant from each site to generate inbred full-sibships (N = 174 sites; 154 from 2017 transplants, plus 20 grown from previous AHQ and RC wild seed collections). Seeds were stratified in 24-well microplates on wet sand for two weeks (4 °C, dark), then germinated at 24°C/ 12°C (1 hr day/12hr night). After 2-3 weeks, seedlings were transplanted to 2” pots (2 pots per maternal family, in randomized positions) and then thinned to one per pot. Plants were maintained under a ~30°C/15°C 16hr/12hr day night cycle with supplemental lighting, bottom-watered to soil-saturation daily, and fertilized monthly. When the first flower on each plant opened fully, we recorded days to flower (1^st^ flower date minus transplant date), 1^st^ flower height (mm) and node number (plus internode length calculated as 1^st^ flower height/node number), and its corolla width, corolla length, and stigma-anther distance (as defined in Fishman et al. 2002). We collected all four anthers of the 1^st^ flower into 50 microL of aniline lactophenol dye and used a haemocytometer to estimate pollen viability and pollen number, following (Fishman and Saunders 2008). At 85 days post-transplant (~3 weeks after the latest 1^st^ flower), we counted the number of stolons and reproductive branches and measured total plant height. Stolons were defined as vegetative branches >1cm long arising below the second leaf pair and remaining horizontal at least to the pot edge, whereas reproductive branches were defined as branches (>1cm long) arising from above the 2^nd^ leaf pair and generally bearing floral buds.

To estimate broadsense heritability (H^2^) and test whether plants from different habitat types (AH vs. PH) differed, on average, in common gardens, we used mixed model REML with habitat as a fixed effect and parental identity as a random effect nested within habitat, implemented in JMP 14 (SAS Institute 2018).

### ddRAD genotyping

Genomic DNA was extracted from bud tissue of the greenhouse-grown wild plants using a CTAB-chloroform protocol modified for 96-well plates (dx.doi.org/10.17504/protocols.io.bgv6jw9e). We used a double-digest restriction-site associated DNA sequencing (ddRADSeq) protocol to generate genome-wide sequence clusters (tags), following the BestRAD library preparation protocol (Ali et al. 2016). Using restriction enzymes *PstI* and *BfaI* (New England Biolabs, Ipswich, MA). An *in silico* digestion of the *M. guttatus* reference genome with this pair generated highly genic and evenly dispersed tags. After enzymatic digestion, sets of 48 individual DNAs were labeled with unique in-line barcoded oligos, and each set pooled into a single tube. Pools were barcoded using NEBNext indexing oligos with a degenerate barcode, PCR amplified, and size-selected (200-500bp fragments) using agarose gels. Libraries were prepared using NEBNext Ultra II library preparation kits for Illumina (New England BioLabs, Ipswich, MA). The libraries were paired-end sequenced (150 bp) on two 1/2 lanes of an Illumina HiSeq4000 sequencer.

Illumina reads were demultiplexed and true PCR duplicates (same restriction site, plus same degenerate barcode) removed using a custom Python script (dx.doi.org/10.17504/protocols.io.bjnbkman). Adapters were trimmed and low-quality reads removed using Trimmomatic (Bolger et al. 2014). Reads were mapped to the *M. guttatus* v2 reference genome (https://phytozome.jgi.doe.gov) using BWA MEM (Li and Durbin 2009) and indexed with SAMtools (Li et al. 2009). We retained 58 million reads after alignment and quality-filtering. After filtering of individuals for mean coverage (>7x) and other quality metrics, we retained genomic data for 231 individuals (N = 119 AH and 112 PH) representing 165 collection sites. SNPs were called using the STACKS pipeline (Catchen et al. 2011; 2013) for aligned sequence data. The “Gstacks” module was used to identify loci and SNPs and to genotype individuals, and we used “Populations” to filter SNPs (maximum missing data = 0.2, minor allele frequency = 0.05) and export genotypes to vcf format. For analyses of population genetic structure and individual genetic diversity, we pruned the dataset to one random SNP per ddRAD tag across the 14 main scaffolds (chromosomes) of the *M. guttatus* v2 reference genome, for a total of 14,919 variable sites. For scans of YNP-wide AH-PH differentiation, we included all variant SNPs (n = 78,479) across those same scaffolds.

### Analyses of population genetic structure and individual variation

Individuals were geo-referenced and named by geyser basin (Fig. 1b), site number within geyser basin, and plant number within site (Supplemental Table S1), but not assigned to populations *a priori*. To explore broad patterns of genetic structure and establish populations for downstream analysis, we used discriminant analysis of principal components (DAPC) implemented in the R package adegenet (Jombart 2008). We used the find.clusters function on the first 20 PCs to identify the number of clusters (k) that minimized the Bayesian Information Content (BIC) and then used the dapc function to cluster individuals (retained PCs = 7, retained axes = 4). To visualize the correlations underlying the DAPC, we performed genetic principal components analysis (PCA) using plink2.0 (Purcell et al. 2007). We also re-calculated the PCA removing all but one individual from the extreme AHQT population. Finally, we used the Bayesian clustering method *conStruct*, which includes continuous geographic distance in model generation, to infer population structure and individual admixture proportions (Bradburd et al. 2018). The three-fold cross-validation method in *conStruct* was used to evaluate k = 2-8 layers, with 3000 Markov-chain Monte Carlo (MCMC) generations assessed for independent convergence for each value of k.

To assess the hierarchical partitioning of genetic variation, we stratified individuals by regions and habitat (excluding AHQT) and conducted analysis of molecular variance (AMOVA) using the R package poppR (Kamvar et al. 2014). Significance of region and habitat effects were tested with the function randtest in package ade4 (Dray and Dufour 2007) using 999 permutations of the data matrix. We also calculated mean pairwise Weir and Cockerham’s F_ST_ (plus bootstrapped 95% confidence intervals) among the nine region x habitat sub-populations using the R program Hierfstat (Goudet 2005). To investigate the landscape scale of genetic differentiation among individuals, we calculated pairwise Euclidean genetic and geographic distance matrices among all individuals (excluding AHQT), as well as generating habitat dissimilarity matrix (AH-AH, AH-PH, and PH-PH pairs of individuals coded as 0, 0.5, and 1, respectively.) We constructed Mantel and partial-Mantel correlograms using mpmcorrelogram (Matesanz et al. 2011) and the ‘vegan’ package in R (Oksanen et al. 2017). Because both geography and habitat had significant effects on genetic distance in the Mantel analyses, we examined the effects of ecology on genetic distance in four geographic distance categories (0-1 km, 1-3 km, 3-10km, >10km) using a model with habitat, distance category, and their interaction and plotted the least square means of the interaction using JMP14.

To examine genome-wide patterns of diversity, we calculated pairwise nucleotide diversity (π) in vcftools (Danecek et al. 2011), for all genotyped individuals, regional populations, and the nine region x habitat subpopulations using variant and invariant sites at 23,545 ddRAD tags (2,409,908 sites) that passed coverage and missing data filters. 95% confidence intervals for mean π values were obtained by bootstrapping. To infer individual inbreeding history, we calculated inbreeding coefficient (F_IS_) using the --het command in vcftools (Danecek et al. 2011), with expected homozygosities calculated within the five regional populations. We compared mean F_IS_ of AH and PH YNP-wide with ANOVA, as well as among regions and all AH-PH partitions within regions. Three individuals with highly negative (< −0.20) values of F_IS_ were excluded from this and the following analyses as potentially contaminated or recent hybrids between distinct populations; this exclusion did not materially affect the results or interpretation. To test for associations between heritable phenotypes (progeny) and inbreeding history (wild plants) within habitat, we excluded all but one AHQT individual, split the dataset by habitat (AH vs. PH) and conducted forward-backward model selection (AICc difference >2 criterion) on all 14 phenotypes as predictors of F_IS_ (traits and F_IS_ standardized to mean and variance = 1). We then examined effects in the best fit models for each habitat using a linear regression framework. F_IS_ statistical analyses were performed in JMP 14 (SAS Institute 2018).

### Tests for independence from widespread annual-perennial inversions

Although dense at >50 tags per Mb or ~10 per cM (~100cM and 15-25 Mb per chromosome; Flagel et al. 2019), our ddRAD dataset does not provide resolution to characterize the genomic basis of geothermal adaptation (Lowry et al. 2016). However, genome scans can rule in or out the involvement of widespread inversions diagnostic of life-history divergence across the species range. Two genomic rearrangements, *inv8* (aka *DIV1*) on Chromosome 8 and *inv5* on Chromosome 5 exhibit elevated F_ST_ in broad-scale annual-perennial comparisons (Oneal et al. 2014; Twyford and Friedman 2015; Gould et al. 2017). *inv8/DIV1* (but not *inv5)* is also a major QTL for numerous traits defining alternative life-history strategies (Hall et al. 2006; Lowry and Willis 2010). To ask whether either inversion was differentiated between AH and AH plants in YNP, we calculated mean F_ST_ in non-overlapping 50-SNP windows across the all-SNP dataset. We then compared mean F_ST_ in windows from genomic scaffolds in *inv8* and *inv5* as defined in (Gould et al. 2017) to the remainder of the genome, using JMP 14.

To independently test whether YNP thermal annuals carry the “perennial” orientation of the widespread *inv8/DIV1* inversion, we estimated local recombination rates in F_2_ test-crosses to the annual (IM; Iron Mountain, Oregon) and perennial (DUN; Oregon Dunes) populations in which *DIV1* was originally identified (Hall et al. 2006; Lowry and Willis 2010). We crossed an AHQT individual (two generations inbred) to both DUN10 and IM106 to make F_1_ hybrids, and then self-pollinated each F_1_ to make F_2_ mapping populations. We genotyped AHQTxDUN10 F_2_s (n = 186) and AHQTxIM106 F_2_s (n = 96) at gene-based markers spanning 1.76 Mb (e173, e278, e675) and 1.85 Mb (e173, e178, e278, e563), respectively, of *DIV1* (see Table S2 for marker details). We expect suppressed recombination in AHQTxIM106 vs. AHQTxDUN hybrids (up to 20 cM difference assuming 5-10 cM/Mb; Flagel et al. 2019) if the YNP annual AHQT carries the widespread perennial version of *inv8/DIV1*, and the opposite if it is chromosomally annual (Lowry and Willis 2010).

## Results

### Heritable variation in life history, plant architecture, and mating system traits

In the greenhouse common garden, plants from thermally-influenced summer-dry habitats (annual; AH) heritably differed from plants from persistently wet habitats (perennial; PH) along life history, plant architecture, and mating system axes (Table 1, Fig. 1c). AH plants produced fewer stolons (4.83 ± 0.65 vs. 13.12 ± 0.76) and more flowering branches (5.56 ± 0.28 vs. 4.15 ± 0.32) than PH plants, resulting in a ~2-fold difference in allocation to sexual vs. asexual reproduction (reproductive ratio: 0.62 ± 0.03 vs. 0.32 ± 0.03). AH plants flowered on earlier nodes with shorter internode lengths, resulting in a much lower height to first flower (Table 1). However, there was only a slight habitat effect on mean days to first flower (AH < 2 days earlier; *P* = 0.03) and no difference in total plant height between AH and PH plants (Table 1). Notably, mean days to first flower was highly variable among AH families (H^2^ = 0.63) and better fit by a mixture of two normal distributions (unimodal normal AICc = 571.2, mean = 39.1, bimodal normal mixture AICc = 544.8, means 37.0 and 44.5) flanking the single intermediate PH peak (mean = 40.8). Broad variation among AH accessions in days to flower under summer daylengths may reflect the diversity of growing season lengths in AH habitats with different thermal influences and water sources (Fig. 1a).

**Table 1.**
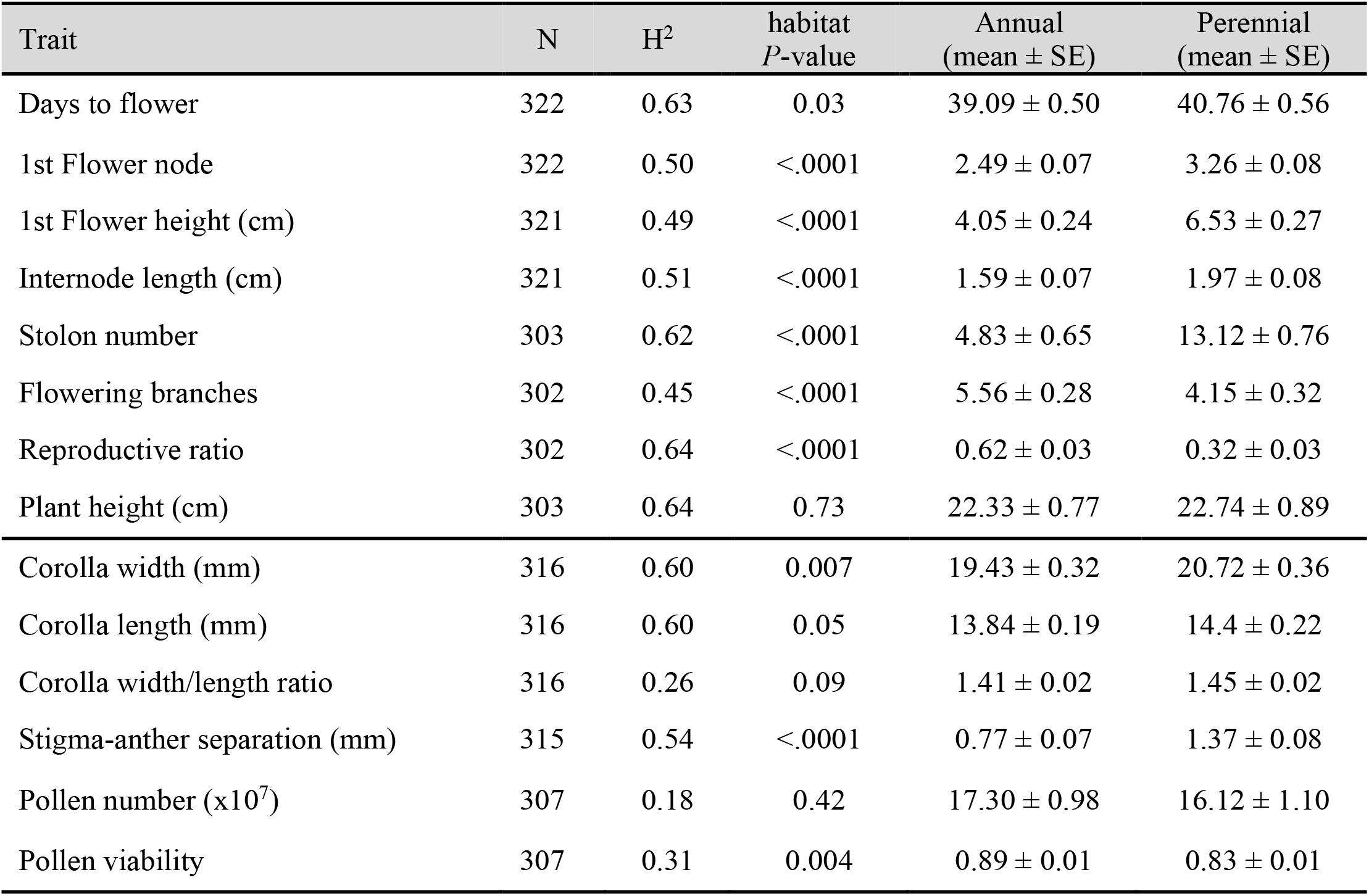
Broad-sense heritability (H^2^) and habitat differentiation of life history and mating system traits in Yellowstone *Mimulus guttatus* grown in a greenhouse common garden. Habitat effects and H^2^ values were derived from mixed model REML analysis with habitat as a fixed effect and full-sib family (selfed, n = 2) as a random effect nested within habitat.

Consistent with selection for autogamous self-pollination and reduced pollinator attraction in spring-flowering annual habitats, AH plants had ~50% lower stigma-anther separation and slightly narrower corollas than PH plants (Fig 1, Table 1). Parallel reductions in corolla length and width-length ratio were only marginally significant (*P* = 0.05 and *P* = 0.09, respectively). Pollen number did not differ among habitats or families, but pollen viability was moderately heritable (H^2^ = 0.38; Table 1) and AH plants had, on average, higher pollen viability than PH plants (0.89 vs. 0.83, P < 0.005; Table 1).

### Population genetic structure across the Yellowstone habitat mosaic

Both discriminant analysis of principal components (DAPC) and *conStruc*t primarily clustered individuals by geography (geyser basin) rather than habitat (Fig. 2). In the DAPC, values of k (clusters) from 5-9 minimized variance among clusters (Fig 2b inset), so we conservatively assigned individuals to five populations (Fig. 2a, b). One consisted entirely of plants from an extreme thermal population AHQT, which was also isolated in previous analyses (Lekberg et al. 2012; Hendrick et al. 2016). The other four clusters correspond to broad geographic regions (designated Central, Northern, Southern, and Southeastern; Fig 2a, b). AHQT’s distinctness does not reflect novel variation; a genomic PCA including only a single AHQT sample places it with Central plants (Fig. S1) and AHQT and the large Central population had similarly low numbers of private alleles per sample (1.3 and 1.1, respectively). *conStruct* also resolved primarily geographical structure (Fig. 2d), and models including a spatial component within layers were always better supported (Fig. S2). At the k=5 supported by cross-validation (Fig. S2), only AHQT individuals were >85% assigned to a single layer and individuals within each DAPC-defined region showed substantial admixture (Fig. 2d). Furthermore, at k=2-8 layers, habitat never defined a discrete component (Fig. S3).

**Fig. 2.**
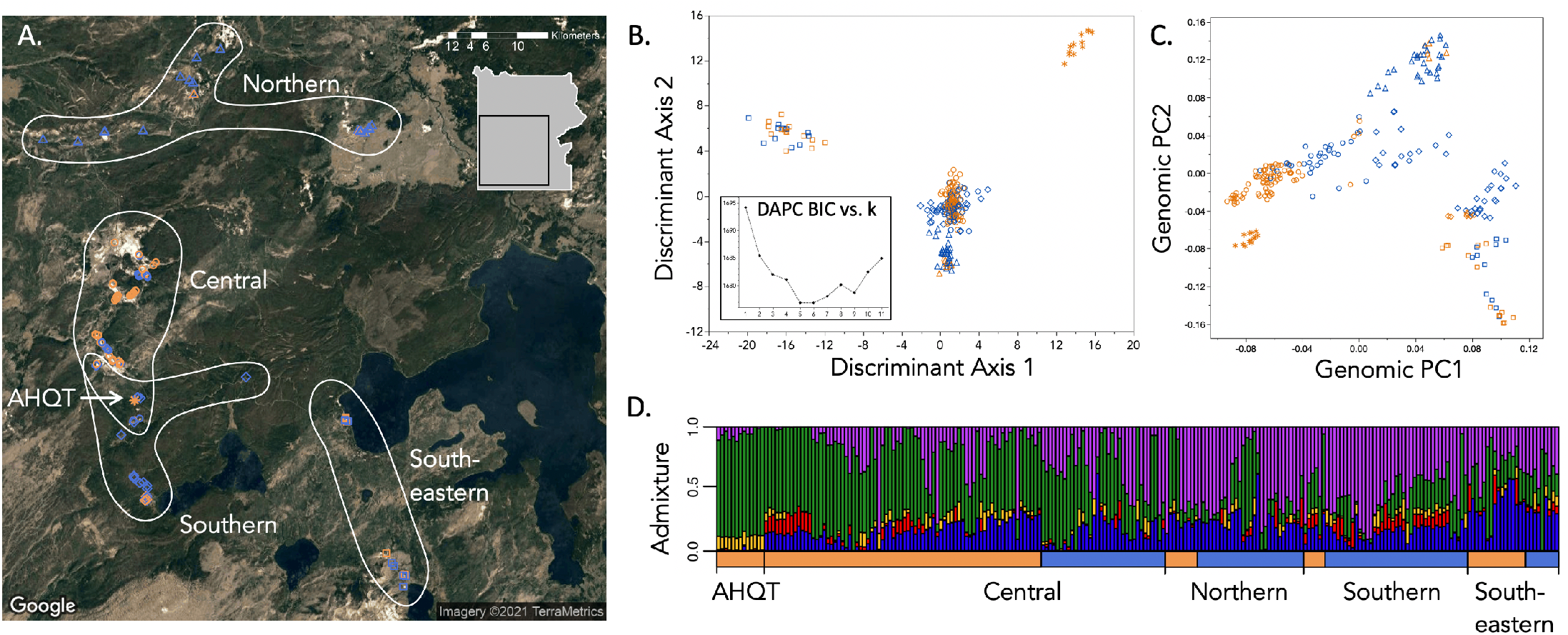
Population genetic structure in Yellowstone *Mimulus guttatus* is primarily geographical, rather than habitat-associated. **(A.)** Individual sample locations coded by habitat (orange = annual, blue = perennial) and population assignments from DAPC (k = 5). The colors and population symbols are repeated in B-D. **(B.)** Individuals plotted along the first 2 discriminant axes of the DAPC with best k = 5. Inset: Bayesian Information Criterion values for DAPC at k clusters 1-11. **(C.)** Individuals plotted along the first two genomic principal component (PC) axes contributing to the DAPC assignments. Including only one individual from AHQT in the PCA shifts it to within the Central cluster, while other PC scores remain very similar (Fig. S1). **(D.)** Admixture proportions of individuals from *conStruct* analysis (k = 5), ordered by habitat and DAPC-defined population.

The four DAPC-defined regional populations were moderately differentiated from each other (F_ST_ = 0.10 - 0.18) and AHQT was highly differentiated from all other populations (F_ST_ = 0.23 - 0.35), but AH and PH partitions within each region were largely undifferentiated (F_ST_ = 0.01 - 0.05) (Table 2). Geographical region (AHQT excluded) explained a much larger portion of the variance (populations; ~12%) than habitat within region (subpopulations; ~2%) in AMOVA, but both components were much smaller than within-subpopulation variation (~28%) (Table 3). However, all AMOVA components were significantly non-zero, consistent with the ddRADs sampling some loci contributing to local adaptation.

**Table 2.** Pairwise F_ST_ (and 95% confidence intervals) among the five DAPC-defined population clusters, including the extreme thermal annual AHQT, and between annual and perennial habitats within each regional population. Confidence intervals were calculated with 100 bootstrapped replicates

| Population pair | $F_{ST}$ | 95% C.I. |
| --- | --- | --- |
| AHQT / Southern | 0.279 | 0.277 - 0.281 |
| AHQT / Central | 0.229 | 0.227 - 0.231 |
| AHQT / Northern | 0.302 | 0.300 - 0.304 |
| AHQT / Southeastern | 0.348 | 0.346 - 0.350 |
| Southern / Central | 0.108 | 0.107 - 0.109 |
| Southern / Northern | 0.101 | 0.100 - 0.102 |
| Southern / Southeastern | 0.119 | 0.118 - 0.120 |
| Central / Northern | 0.124 | 0.122 - 0.125 |
| Central / Southeastern | 0.170 | 0.168 - 0.172 |
| Northern / Southeastern | 0.182 | 0.181 - 0.184 |
| AH / PH Southern | 0.049 | 0.047 - 0.050 |
| AH / PH Central | 0.027 | 0.026 - 0.027 |
| AH / PH Northern | 0.013 | 0.012 - 0.014 |
| AH / PH Southeastern | 0.012 | 0.011 - 0.013 |

**Table 3.** Analysis of molecular variance (AMOVA) across DAPC-assigned regions and habitats (AH v. PH). PVE is the Percent of Variance Explained by each hierarchical factor, and significance (*P*) vs. zero was calculated by random permutation (n = 999) of the data matrix.

| Variance component | df | PVE | $P$ |
| --- | --- | --- | --- |
| Among regions (excluding AHQT) | 3 | 11.8 | < 0.001 |
| Between habitats within regions | 4 | 2.3 | < 0.001 |
| Among individuals within region/habitat | 210 | 27.6 | < 0.001 |

Mantel correlations indicated substantial isolation-by-distance and moderate isolation-by-habitat (Mantel: geography *r* = 0.56; habitat *r* = 0.16; both P <0.0001; partial Mantel values were near identical). Notably, the relationship between genetic and geographic distance depended on habitat at local spatial scales, with PH-PH pairs more genetically similar than AH-PH and AH-AH pairs at distances < 1km but AH-AH pairs more similar at greater distances (Fig. 3).

**Fig. 3.**
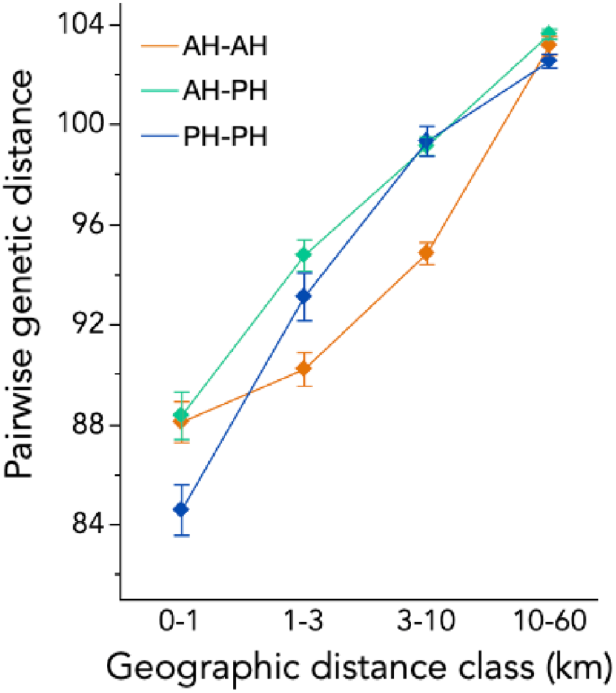
Mean (± SE) pairwise genetic distance for pairs of individuals sampled from annual, annual-perennial (mixed), and perennial habitats across geographical distance categories.

### Habitat and phenotypic effects on genetic diversity

YNP *M. guttatus* exhibited moderate nucleotide diversity (π = 0.011), higher than genome-wide genotyping-by-sequencing estimates for sparsely-sampled perennial *M. guttatus* populations globally (all π < 0.01;Vallejo Marín et al. 2021) but similar to one deeply sequenced annual population (π = 0.014; Puzey et al. 2017). Regional populations captured similar diversity (π = 0.009-0.011), as did AH (π = 0.008-0.010) and PH subsets (π = 0.009-0.011) within each region. AHQT was about half as diverse as other partitions (π = 0.005) and relatively invariant at sites segregating YNP-wide (55% of sites polymorphic vs. 87-98% for the regional mixed populations). Despite its lower expected heterozygosity, the mean inbreeding coefficient (F_IS_) at AHQT was significantly higher than all other regional populations (Table 4). AH plants overall (excluding AHQT) also had significantly higher F_IS_ (mean = 0.41 ± 0.02 SE, N = 103) relative to PH individuals (mean = 0.29 ± 0.02 SE, N = 110; habitat r^2^ = 0.065, P < 0.0001; Fig. 3; Table 4). This pattern was driven by the larger majority-annual Southeastern and Central populations; in the majority-perennial Northern and Southern regions, AH did not have greater F_IS_ (Table 4).

**Table 4.** Mean nucleotide diversity (π) and inbreeding coefficients (F_IS_) ± SE (plus n for both analyses) of *M. guttatus* plants sampled from annual (AH) and perennial (PH) habitats in DAPC-defined regional populations in Yellowstone National Park. For all estimates of π, the 95% bootstrapped confidence intervals were within the digits shown for the means.

| Population | $\pi$ (AH) | $\pi$ (PH) | $F_{IS}$ (AH) | $F_{IS}$ (PH) |
| --- | --- | --- | --- | --- |
| AHQT | 0.005 | n/a | $0.52 \pm 0.04$ (13) | n/a |
| Central | 0.010 | 0.010 | $0.42 \pm 0.03$ (78) | $0.27 \pm 0.04$ (33) |
| Northern | 0.009 | 0.011 | $0.22 \pm 0.07$ (6) | $0.38 \pm 0.04$ (29) |
| Southern | 0.008 | 0.011 | $0.25 \pm 0.05$ (7) | $0.25 \pm 0.02$ (39) |
| Southeastern | 0.009 | 0.009 | $0.43 \pm 0.09$ (13) | $0.25 \pm 0.05$ (11) |
| All | 0.011 | | $0.40 \pm 0.02$ (117) | $0.28 \pm 0.02$ (111) |

AH plants (even excluding AHQT) exhibited bimodality of F_IS_ values (Fig. 4) and parental inbreeding coefficients were correlated with heritable traits. The AH F_IS_ model with minimum AICc (r^2^ = 0.48) included height to first flower, corolla width/length ratio, stigma-anther separation, and pollen viability (Table 5; N = 40 families with all traits measured). Higher F_IS_ values were associated with flowering closer to the ground, reduced herkogamy, relative wide corollas, and lower pollen viability. Among PH plants (n = 44), no phenotypic variables improved prediction of F_IS_ over the null model.

**Table 5.** Parental inbreeding coefficient (F_IS_) prediction by common-garden progeny phenotypes across annual habitat (AH) *M. guttatus* lineages (N = 40 with genome-wide F_IS_ values and all traits measured in multiple progeny). The best-fit model (r^2^ = 0.48) was obtained by forward-backward stepwise model selection using a minimum AICc criterion (AICc for this model = 102.4 vs. 107.3 for 3-trait model, 101.3 for 5-trait model, and 118.1 for null model).

| Trait | $F_{IS}$<br>Effect<br>Estimate | $F_{IS}$<br>Effect<br>SE | $F_{IS}$<br>Effect<br>P-value |
| --- | --- | --- | --- |
| 1<sup>st</sup> flower height | -0.51 | 0.14 | 0.001 |
| Corolla W/L ratio | 0.38 | 0.12 | 0.005 |
| Stigma-anther separation | -0.30 | 0.12 | 0.023 |
| Pollen viability | -0.29 | 0.13 | 0.031 |

**Fig. 4.**
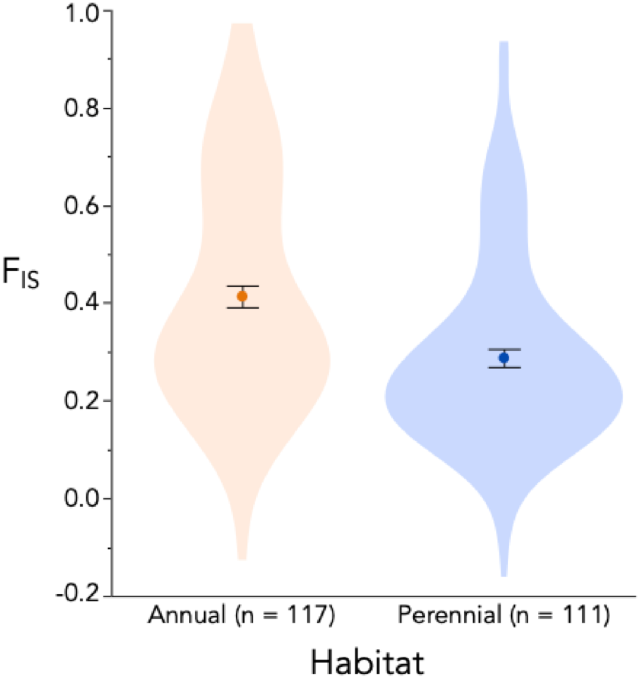
Means, standard errors, and distributions of individual inbreeding coefficients (F_IS_) of *M. guttatus* plants sampled from annual and perennial habits in Yellowstone National Park. Individual F_IS_ values were calculated within each the five regional populations. The significant habitat effect (p < 0.001) and bimodality of the annual samples persists if all AHQT individuals are removed.

### Tests for genetic independence of annuality in Yellowstone M. guttatus from widespread inversions

The *DIV1/inv8* inversion, which is phenotypically and genetically diagnostic of life history elsewhere in the species range, did not show elevated F_ST_ between AH and PH plants in Yellowstone (Fig. 5a). Indeed, sites in *DIV1* had slightly reduced differentiation relative to the rest of the genome (mean F_ST_ = 0.02 vs. 0.03, Fig. 5b). However, the inv5 region exhibited substantially elevated F_ST_ (mean = 0.09; Fig. 5b) (as it does range-wide between annuals and perennials despite no phenotypic effects on life history; Gould et al. 2017), and contained 25% (10/40) of the top 2.5% outlier windows.

**Fig. 5.**
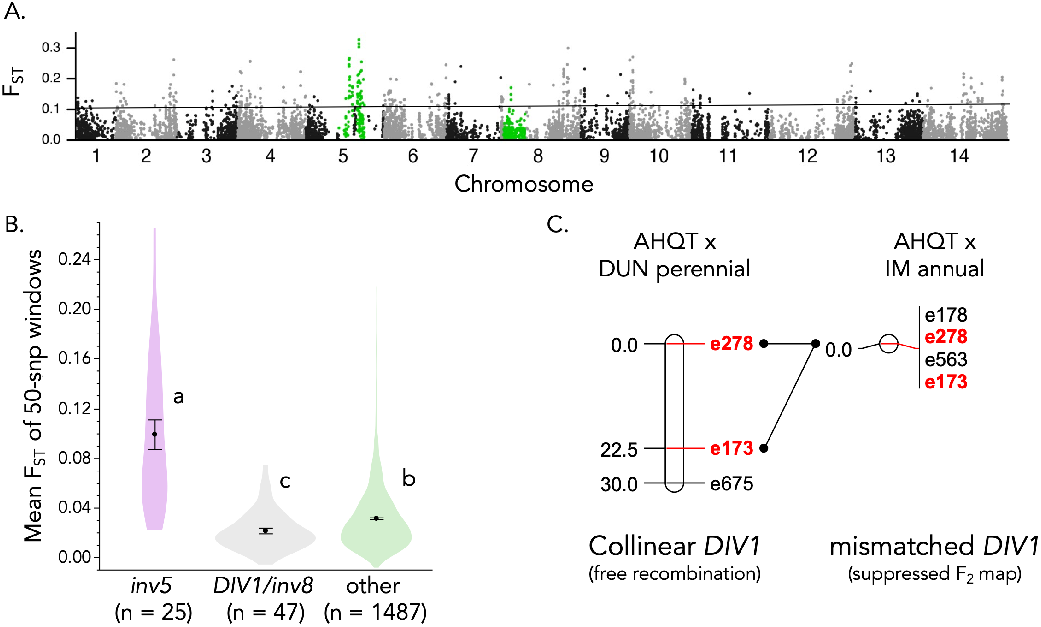
Genome scan and linkage tests indicate the independent derivation of annuality in Yellowstone *M. guttatus*. **(A.)** F_ST_ between AH and PH plants at SNPs genome-wide. Highlighted SNPs are in inversions on Chromosomes 8 and 5 that phenotypically define annuals and perennials (*inv8/DIV1* only; Hall et al. 2006; Lowry and Willis 2010) or show elevated annual-perennial F_ST_ range-wide (both; Gould et al. 2017). **(B.)** Means, standard errors, and distributions of mean F_ST_ for 50-SNP windows genome-wide. Letters indicate means significantly different by Tukey’s HSD test. **(C.)** Linkage test for orientation of the *DIV1* region in AHQT Yellowstone annual, with free recombination (>20 cM) in hybrids with coastal perennial DUN and suppression of recombination (inversion) in hybrids with IM106 Oregon annual indicating AHQT collinearity with the widespread perennial haplotype.

Targeted linkage mapping using a thermal annual (AHQT) test-crossed to lines with the annual (IM106) and perennial (DUN) arrangements of *DIV1* further demonstrated that thermal annual *M. guttatus* are “perennial’ at this widespread diagnostic locus. Across ~2Mb of *DIV1*, recombination was strongly suppressed in AHQTxIM106 hybrids relative to AHQTxDUN10 hybrids (Fig. 5c) indicating that YNP annuals are collinear with widespread (and likely local) perennials. Together, these lines of evidence indicate no re-use of the widespread *DIV1/inv8* chromosomal polymorphism in Yellowstone life-history adaptation and suggest that phenotypic differentiation between YNP thermal annuals vs. nonthermal perennials arose independently.

## Discussion

### Life history and mating system evolution in Yellowstone thermal annuals independently parallels range-wide patterns of M. guttatus adaptation to ephemeral habitats

Our results demonstrate the power of natural selection to generate complex reproductive and edaphic adaptation despite abundant gene flow across a microgeographic habitat mosaic. Although based on only a single observation of late summer dry-down at each site, the definition of annual (AH) vs. perennial (PH) sites captured the major axes of phenotypic adaptation in Yellowstone monkeyflowers remarkably well (Fig. 1c, Table 2). These habitats are strongly shaped by geothermal influences on YNP soils (i.e., all AH habitats are in active thermal areas), but soil temperature *per se* is likely not the major selective agent. Instead, like widespread *M. guttatus* ecotypes (Hall and Willis 2006; Lowry and Willis 2010) and *M. cardinalis* from Southern California, geothermal annual plants have adapted via drought-escape into an alternative temporal niche for growth and reproduction. The resulting phenotypic differentiation parallels broader-scale patterns in *Mimulus*, but with YNP-specific features that illuminate general rules governing life history and mating system evolution.

Stolons (or rhizomes) allow for overwintering after survival through summer, and their loss is diagnostic of drought-associated annualization in *Mimulu*s (Hall and Willis 2006; Lowry and Willis 2010; Nelson et al. 2021) and other herbaceous plants (reviewed in Friedman and Rubin 2015; Friedman 2020). Yellowstone thermal AH plants were genetically annualized, producing few stolons, more reproductive branches, and a 2-fold higher ratio of sexual to asexual reproduction (Table 1, Fig. 1c). Negative correlations between stolons and flowering shoots (Fig. 1c) may reflect either a genetic tradeoff in meristem allocation or correlated selection across the landscape (Friedman and Rubin 2015; Friedman 2020). Interestingly, increased AH floral allocation does not consistently result from the shift to earlier flowering often seen in annuals (Friedman et al. 2015); flowering time under summer-like conditions showed high heritability and significant bimodality in annuals, but only weak AH-PH differentiation (Table 1). This contrasts with the widespread life history polymorphism in *M. guttatus*, where annuals exhibit a “fast” strategy characterized by rapid flowering (Kooyers et al. 2015; Troth et al. 2018) and stolon number and flowering time are positively genetically correlated (Lowry and Willis 2010; Friedman et al. 2015). AH plants in highly thermal sites may have long growing seasons (e.g., November to June) relative to both PH plants and AH plants where soils are not hot enough to remain snow-free all winter. This suggests that finely-tuned natural selection can readily break-up expected correlations among life history traits; however, additional work on phenological variation will be necessary to understand whether and how multiple traits contributing to life history strategies are packaged in YNP.

Strong reductions in stigma-anther separation (aka herkogamy) in AH plants (Fig. 1, Table 1) likely promote autogamous self-pollination (reviewed in Sicard and Lenhard 2011; Opedal 2018), providing reproductive assurance during spring flowering prior to pollinator emergence (Lekberg et al. 2012). Consistently low stigma-anther separation in AH flowers is consistent with natural selection (Fishman and Willis 2008) and experimental evolution (Bodbyl Roels and Kelly 2011) in the absence of pollinators in *M. guttatus*, and with the generally high evolvability of herkogamy across plant populations (Opedal et al. 2017). Derived selfer species of the species complex (e.g. *M. nasutus*) show a similar association with ephemeral habitats, even relative to sympatric *M. guttatus* annuals (Martin and Willis 2007; Ferris et al. 2014; Kenney and Sweigart 2016). However, AH plants likely retain opportunities for pollinator attraction and outcrossing, as even the smallest AH corollas were absolutely larger than many highly outcrossing annual *M. guttatus* (Bodbyl Roels and Kelly 2011). Reductions in corolla size associated with the “selfing syndrome” may arise through two non-exclusive mechanisms (Sicard and Lenhard 2011): adaptive shifts in energy investment after selfing becomes routine (Charlesworth & Charlesworth 1981) and/or fixation of formerly deleterious recessive “small” alleles by drift (Fishman et al. 2002). Plants in extreme thermal annual sites (e.g., AHQT) may flower and die early enough to rarely encounter pollinators, promoting corolla reduction via all mechanisms and putting them on the path toward obligate selfing (Lekberg et al. 2012). However, AH individuals in less extreme locations (e.g., Shoshone Geyser Basin, where both AH and PH plants were flowering in late July) may continue to interact with pollinators and exchange genes with PH mates later in their phenology, and so maintain large corollas through both selection and gene flow.

Variation among YNP AH plants provides a unique opportunity to understand the consequences of mating system evolution in its very early stages. Our estimates of F_IS_ suggest abundant and structured variation --individual inbreeding coefficients were shaped by habitat (Fig. 4) and (in AH plants) associated with heritable phenotypes for four traits (Table 5). The negative correlation of stigma-anther separation with parental inbreeding coefficient in YNP annuals supports herkogamy’s importance to autogamous self-pollination (Opedal 2018). Similarly, the strong predictive power of first flower height is likely a proxy for extreme phenology, as short stature adapts plants to the narrow boundary layer of winter-warm air typical of the most extreme thermal sites (Lekberg et al. 2012). However, the positive association of corolla width/length ratio with F_IS_ is unexpected, as relatively narrow corollas were favored by selection for efficient autogamy in annual Oregon *M. guttatus* (Fishman and Willis 2008) and focal AH populations had narrower flowers than PH populations (Lekberg et al. 2012). This difference may reflect inbreeding depression of corolla width in this study or may possibly be an artifact of population structure. Higher pollen viability of experimentally inbred AH vs. PH plants (Table 1), as well as the negative correlation between pollen viability and F_IS_ within AH plants (Table 5), also suggests variation in genetic load among habitats and individuals with distinct mating histories. However, because AH-PH differentiation largely matches previous (outbred) common gardens (Lekberg et al. 2012), inbreeding depression is unlikely to account for the overall phenotypic patterns.

The phenotypic diversity of thermal annual *M. guttatus* (Fig. 1) may in part arise from the distinct genetic basis of their life history shift, which is independent of the widespread *DIV/inv8* inversion polymorphism (Fig 4). *DIV1* controls numerous traits, including stem thickness, corolla size, herbivore defense and flowering time, as well as stolon production, range-wide (Hall et al. 2010; Lowry and Willis 2010; Lowry et al. 2019). By packaging co-favored variation into alternative haplotypes spanning 100s genes, inversions can consolidate and accelerate local adaptation in the face of gene flow (Kirkpatrick and Barton 2006). However, suppressed recombination in heterozygotes for established alternative haplotypes (as with *DIV1*) may also limit genetic combinations and constrain adaptation to novel habitats. As yet, we cannot say why the widespread *DIV1* inversion polymorphism was not used to adapt to novel annual habitats in YNP or what genomic regions account for the convergent evolution of annuality there. However, Yellowstone annuals are clearly built on perennial *DIV1* chromosomal chassis (Fig. 5), and thus key life-history traits (e.g., stolon number) evolved independently from both genetic and ecological perspectives. Unlike *DIV1*, *inv5* has not been associated with life-history traits in QTL mapping (Hall et al. 2006; Lowry and Willis 2010; Friedman 2014; Friedman et al. 2015), but elevated F_ST_ in YNP mirrors widespread annual-perennial differentiation in this region (Gould et al. 2017). The only known phenotypic association of *inv5* region is with vernalization requirement in hybrids between inland and coastal perennials (Friedman and Willis 2013), suggesting it may also segregate within perennials. Selection on similar phenological cues, and/or below-ground or physiological traits, may explain YNP differentiation across *inv5*. Finally, because our ddRAD scan did not reveal extended F_ST_ elevation elsewhere in the genome (Fig. 5), correlated shifts in life history, plant architecture, and reproductive traits are likely not coordinated by a novel large inversion (i.e., one >5 Mb like *inv5* and *DIV1*).

### A little gene flow goes a long way: habitat and phenotypic variation influence individual mating patterns, but only lead to substantial AH-PH reproductive isolation in micro-allopatry

Patterns of genome-wide genetic variation in YNP M. guttatus illustrate three distinct outcomes of strong natural selection on life-history and reproductive traits across a complex habitat matrix. First, genetic variation was largely structured by geography rather than ecology, with AH-PH adaptation occurring on a background of abundant genetic exchange within multiple regions. Second, despite the general lack of habitat-associated population structure, local adaptation and direct environmental effects generate habitat-specific differences in individual inbreeding coefficient and isolation by distance. Third, one large thermal annual population (AHQT), which is both ecologically extreme and geographically isolated by a ~200m gap of no *M. guttatus* habitat, exhibits the signatures of incipient (but still reversible) speciation. These population genetic outcomes of life-history and mating system divergence are a microcosm of those seen across the entire *M. guttatus* species complex, as well as broadly across plant species. Manifested over short spatial (and likely temporal) scales in this unique system, these results provide a window into common evolutionary processes often confounded by geography and multiple layers of reproductive isolation.

Previous work in Yellowstone *M. guttatus*, focused on two sites with adjacent thermal annual and nonthermal perennial extremes, found high genome-wide differentiation in one (F_ST_ = 0.2-0.3 at AHQ; (Lekberg et al. 2012; Hendrick et al. 2016) and negligible habitat-associated structure in the other (Lekberg et al. 2012). Here, ddRAD genotyping across all major thermal basins demonstrates predominance of the latter pattern, indicating that shared AH-PH phenotypic divergence generally represents adaptation in the face of gene flow. Multiple clustering approaches (Fig. 2) and estimation of the effects of habitat within the context of geographical structure (Tables 2 and 3), found little genome-wide differentiation between AH-PH plants. This parallels the predominantly geographical structure of widespread annual and perennial *M. guttatus* across the species range (Oneal et al. 2014; Twyford and Friedman 2015; Gould et al. 2017).

Adaptation with gene flow is common, but microgeographic edaphic mosaics like YNP require strong natural selection to shape multi-trait adaptive strategies. Theory predicts that major (and antagonistically pleiotropic) loci should underlie the phenotypic differentiation resistant to gene flow in the absence of extrinsic barriers (Yeaman and Whitlock 2011). For Mendelian traits, like pigmentation (e.g., Jones et al. 2018) or copper tolerance (e.g., Lee and Coop 2017), such phenotypic sorting by strong selection may generate only low migration load. However, maintaining polygenic multi-trait syndromes (such as life-history strategies) in the face of gene flow may generate high load or local maladaptation. Thus, as theory predicts (Kirkpatrick and Barton 2006), complex alternative strategies are often associated with chromosomal inversions, which suppress recombination (in heterozygotes) across large genomic regions containing many functionally diverse genes (Tigano and Friesen 2016; reviewed in Mérot et al. 2020). Here, while we can rule out re-use of the functionally important *DIV1/inv8* inversion, the genetic basis of correlated trait divergence (Fig. 1c) remains an open question. However, background genetic homogenization between YNP annual and perennials will provide an ideal framework for exploring the genetic architecture of complex adaptation in the face of gene flow.

Although the overall population structure of YNP *M. guttatus* is geographic, reproductive adaptation to unique temporal niche created by thermal AH habitats nonetheless leaves genomic signatures. Theory predicts that shifts in mating system profoundly affect patterns of individual genetic variation, as well as population genetic diversity and connectivity (reviewed in Wright et al. 2008; 2013; Hu 2015). Over the short term, an abrupt shift to exclusive self-fertilization (e.g., loss of pollinators) sorts existing variation among increasingly homozygous individuals. However, in the many plant populations or species that have evolved the floral selfing syndrome, genetic diversity is further minimized by low effective population size, an ecological propensity for bottlenecks, and accentuated background selection (St Onge et al. 2011; Barrett et al. 2014; Barrett and Harder 2017; López-Villalobos and Eckert 2018). In Yellowstone, we see something closer to the expectation for a recent shift in mating system. Each AH “sub-population” carries nearly the same genetic variation (both in identity and quantity) as the PH plants in the same geographical region, but that variation is (on average) more homozygous in AH individuals (Fig. 3). Denser sampling (<100m scale) might reveal small sub-populations of related AH plants within thermal patches, but AH plants were relatively isolated from each other and PH plants nearby (<1km), while AH-PH and PH-PH pairs were relatively isolated at intermediate distances (1-30km) (Fig. 4). Habitat differences in the slope of isolation-by-distance may reflect clonality in PH plants as well as AH selfing, but it persisted when the 10 most close genetic pairs (possible clones; 9 PH, 1 AH, all within 10m) were removed (data not shown). The key role of selfing is underlined by our observation that F_IS_ in wild plants was correlated with both habitat (Fig. 3, Table 3) and (in AH plants) by traits suggestive of the local strength of selection for selfing and spring flowering (Table 5). Finer-scale sampling and direct measures of outcrossing rates (e.g. Colicchio et al. 2020) and inbreeding history will be necessary to tease apart the contributions of the temporal/spatial environment (including pollinators and mates) and reproductive phenotypes on patterns of mating and genetic relatedness across the mosaic of YNP *M. guttatus* habitats. However, preliminary evidence that individual inbreeding history is phenotypically predictable suggests novel opportunities for investigating the early dynamics of mating system evolution, including the lineage-specific evolution of inbreeding depression.

Overall, our results indicate that a multi-trait adaptive syndrome, including floral traits allowing 100% autogamous self-pollination (Lekberg et al. 2012), has evolved within a longer-term context of regional genetic exchange. Even the most extreme AH individuals appear to be built from essentially random subsets of local PH variation, plus alleles at a limited number of loci under selection. Given the spatial and temporal patchiness of the YNP geothermal matrix (Fig. 1, Fig 2), AH-adapted alleles likely have an extensive history of transiting through relatively contiguous PH populations, and then re-assembling (with help from selection as well as inbreeding) in distant or newly-erupted thermal annual sites. This might resemble, on a vastly smaller spatial and temporal scale, the global dynamics of sticklebacks, where (to a large extent) local adaptation to newly appearing freshwater habitats occurs from standing variation persisting as rare maladaptive variants in the marine environment (Colosimo et al. 2005; Nelson and Cresko 2018). This scenario strongly contrasts with the idea that mating system and flowering phenology act as “magic traits” whose divergence can generate sufficient reproductive isolation to initiate plant speciation in sympatry (e.g. Savolainen et al. 2006). Here, despite shifts in mating system and phenological traits sufficient to generate predictable habitat differences in individual outcrossing rate and connectivity, we do not see the build-up of population-level reproductive isolation. Although definitions of species vary (Harrison 2012), this is the exact opposite of speciation (i.e., the maintenance of genome-wide genetic continuity despite strong natural selection on reproductively isolating traits).

The one exception to the overall pattern, AHQT, provides a second window into evolutionary outcomes of strong microgeographic selection affecting reproductive traits. Although plants from this population share a common thermal annual phenotype with other AH plants (few/no stolons, low height at first flower, zero stigma-anther separation), they are highly differentiated genome-wide from all other populations (F_ST_ ~ 0.2) and showed reduced π, polymorphism, and heterozygosity despite being sampled over a similar spatial scale. The distinctness of AHQT in part reflects the uniform ecology of this large thermal site, which is geothermally extreme (soils > 60°C in summer), has no source of soil moisture other than snow, and is particularly early flowering (Lekberg et al. 2012). However, in addition to these factors, a key trigger for its high differentiation is likely a ~200m gap of no *M. guttatus* habitat separating all of AHQT from the nearest (late-flowering perennial) AHQN population. This discrete gap acts as a geographical barrier to gene flow, facilitating both unimpeded thermal-soil specialization (including in traits that increase isolation) and the neutral differentiation by drift (including bottlenecks) typical of established selfing populations and species. This perfect storm of selection, drift via high selfing rates, and very low migration likely recapitulates the early stages of speciation in selfer taxa like *M. nasutus*, which represents a small, highly homozygous, and geographically-specific subsample of *M. guttatus* variation despite its current broad range (Brandvain et al. 2014). Alone, each factor would likely be insufficient to fully isolate AHQT over such a short spatial scale, but together they appear to allow the first steps toward speciation. Thus, while “magic” trait divergence undoubtedly contributes to the evolution of reproductive isolation in most geographical contexts, a boost from space (plus time) may be necessary for barriers to develop to the point that isolation can persist in full sympatry. In *M. nasutus*, which self-pollinates its tiny, pollen-poor flowers prior to corolla opening (which often doesn’t even occur), that point was likely reached before its spread into extensive range overlap with *M. guttatus*. At AHQT, speciation along a similar path is also conceivable, but could still be undone by the emergence of a thermal stream lined with intermediate AH plants that overlap in flowering time with both extremes. AHQT’s partial isolation, on a continuum with admixture across the rest of the mosaic, suggests that both habitat discontinuity and reproductive adaptation may be necessary to effectively shut down gene flow genome-wide.

## Conclusions

The yellow monkeyflowers of Yellowstone National Park (YNP) exhibit dramatic life history, growth form, and reproductive trait adaptation to the novel edaphic environments created by geothermal activity. This microgeographic divergence parallels differentiation between widespread annual and perennial ecotypes found range-wide, but has a novel genetic basis and reflects the particular selective pressures of the unique YNP thermal habitat(s). With the exception of the extreme (and micro-allopatric) AHQT population, thermal annual plants maintain multi-trait local adaptation in the face of abundant gene flow across the complex habitat matrix, with geography the primary factor structuring genome-wide sequence variation. However, while plants from annual thermal habitats generally sample the same pool of regional variation as their perennial neighbors, their individual genetic variation reveals elevated inbreeding and isolation from each other. Such individual shifts in realized mating system, which integrate floral and phenological adaptation with direct environmental effects, provide rare insight into within-population mating system evolution (particularly in contrast to AHQT and established *Mimulus* selfers). Together, these findings have important implications for understanding the (parallel) evolution of trait syndromes, the structuring of genetic variation by intrinsic and extrinsic factors, and the limits to sympatric speciation.

## Acknowledgments

We are grateful to H. Anderson, A. Carlson, and other Yellowstone National Park staff for facilitating research in Yellowstone, which was conducted under permits YELL-2011/2017/2018-SCI-5834. We thank P. Breigenzer, F. R. Finseth, R. Hanes, M. McIntosh, S. R. Miller, C. L. Pierpont, A. Lapsansky, and M. Hendrick for assistance with field, greenhouse, and/or lab research, and D. Xing of the University of Montana Genomics Core Facility for material support with genotyping and genomics. The research was supported by NSF grants NSF DEB-1457763 and NSF OIA-1736249 to L.F. The pre-print was formatted based on a template: https://github.com/finkelsteinlab/BioRxiv-Template

## Author Contributions

K. M. K. conducted most of the field, greenhouse, and lab work, as well as the bioinformatic analyses, as part of his M.S. thesis under the supervision of L.F. C. B. performed some of the population genetic analyses and assisted with data curation. T.C.N. developed genomics protocols and assisted with lab work and bioinformatics. L.F. conceived of the research, assisted with field, greenhouse, and lab work, conducted phenotypic and linkage analyses, and constructed most figures. The manuscript was primarily written by L.F. and K. M. K., with contributions and editing from all authors.

## Data Accessibility

Genomic data will be posted on the NCBI Sequence Read Archive under Bioproject PRJNA744896, and the phenotypic and derived genetic datasets will be archived on Dryad at publication.

*The authors declare no conflict of interest*.

