## Supplemental Tables & Figures for "Population genomic consequences of novel life history and mating system adaptation to a geothermal soil mosaic in yellow monkeyflowers (*Mimulus guttatus*)"

### Supplemental Tables and Figures

**Table S1.** See Excel sheet with sample information.

**Table S2.** Markers used for *DIV/inv8* recombination test.

| MgSTS marker | V2 sc_8 position | V1 scaffold | F primer | R primer |
|---|---|---|---|---|
| e675 | 2044482 | 11 | TCGTTGGTGGAAATCAAAGC | CCGGATCCTTGTTGAGTAGC |
| e173 | 2444182 | 11 | GGCAATCTTCAACCTTTTCC | AGGCTGGTGGCTTATCTTCC |
| e278 | 3605725 | 11 | ACGTCAGCCCTTTGTACACC | ACTCAGTTGTGCCAGTCACC |
| e563 | 3695076 | 11 | AAACTGGTGTGCTGAAAGAGC | GCTTCCACCGTAAATTCTCC |
| e178 | 6759231 | 233 | CCTCGTTGTCTTTGTTCAGG | TGAAACGCCAGTTTGATGC |

Figure S1. The first two principal components of genomic variation for YNP *M. guttatus* individuals, coded by habitat (AH = orange, PH = blue) and DAPC-defined region (circle = Central, diamond = Southern, triangle = Northern, square = Southeastern), calculated with only a single AHQT individual (star) in the dataset.

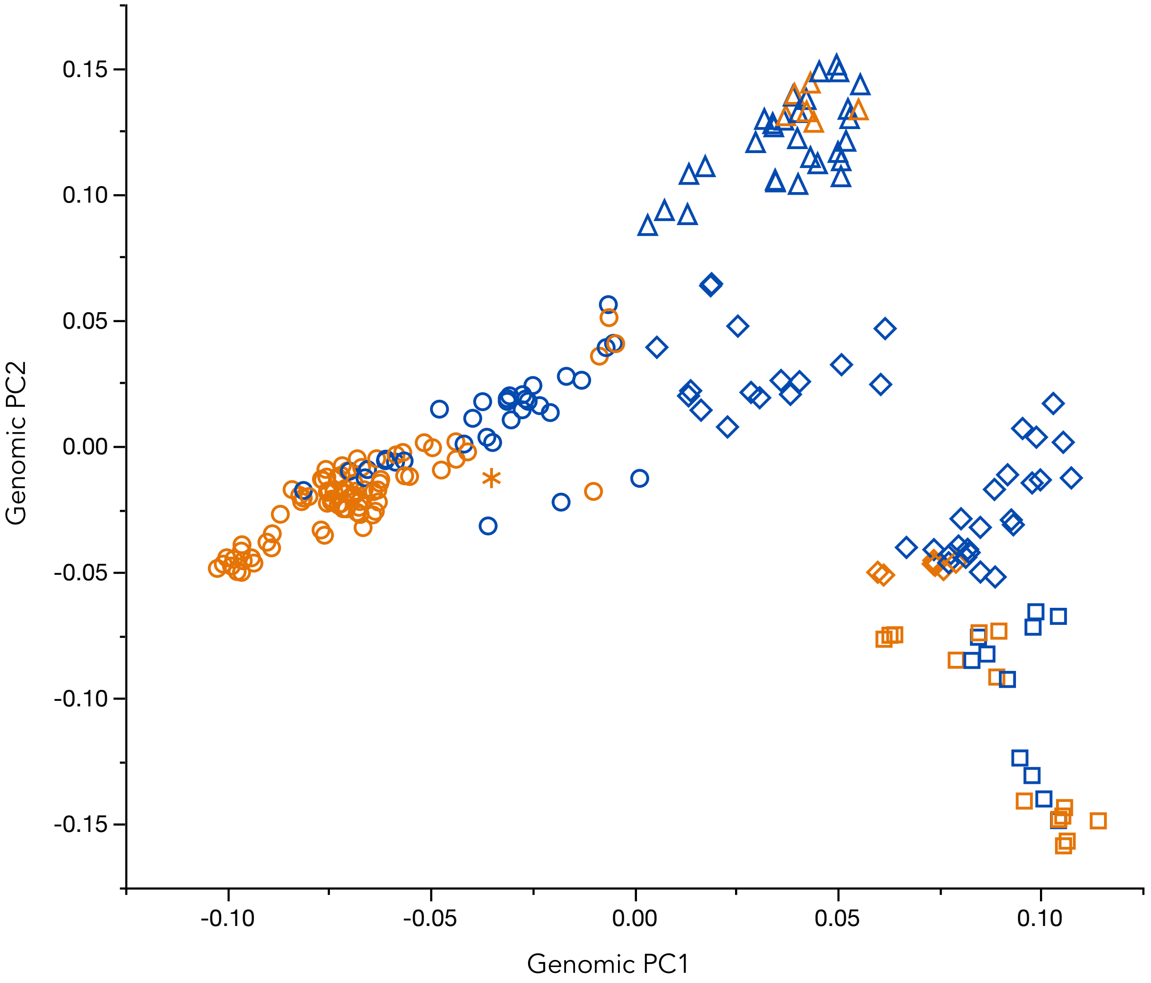

Figure S2. Cross-validation plots for *conStruct* analyses. A. Predictive accuracy of spatial (blue) and non-spatial (green) models across values of k. B. Close-up of the spatial cross-validation results, showing asymptotic accuracy at k = 5 (bars show 95% confidence intervals).

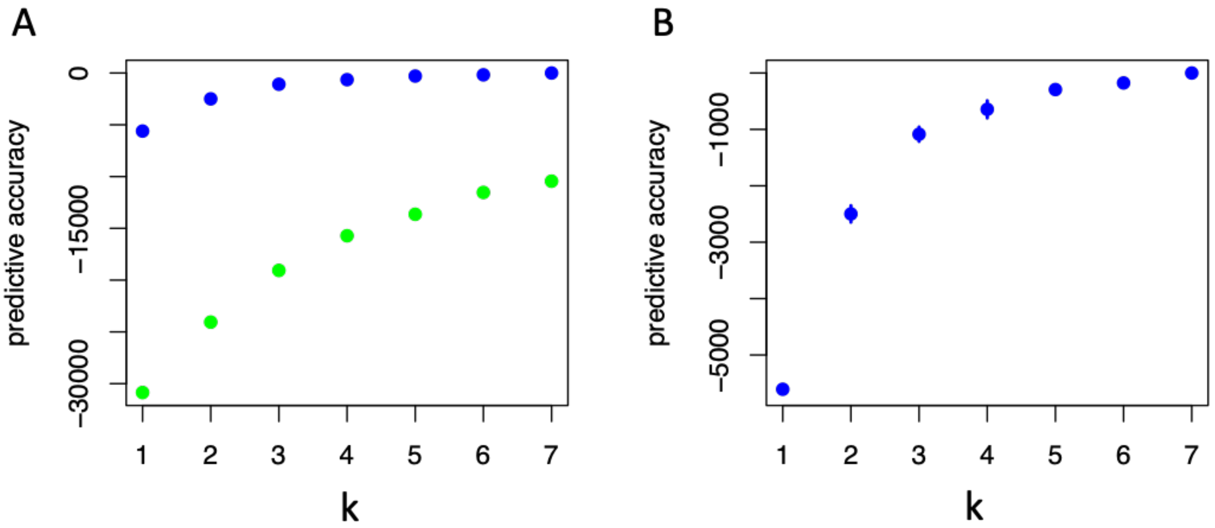

**Figure S3.** Admixture proportions from *conStruct* with k = 2-4 and 6-8 layers. Individuals (bars) are ordered by habitat (AH = orange, PH = blue) and DAPC region as in Fig. 2d (showing k = 5).

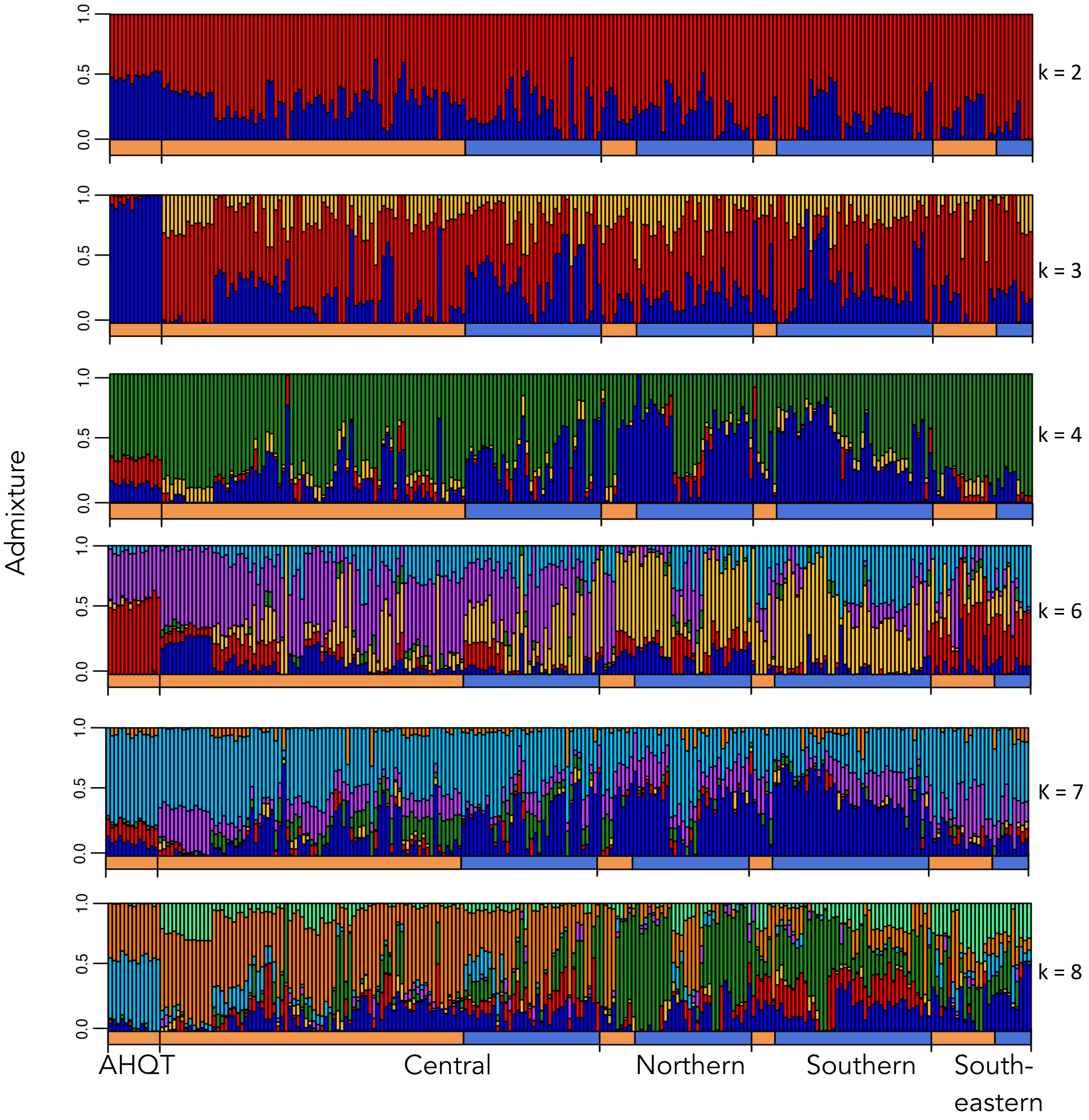
