## Supplementary material for "Population genomic consequences of novel life history and mating system adaptation to a geothermal soil mosaic in yellow monkeyflowers (*Mimulus guttatus*)": Table S1

| Individual | Dataset | Habitat | Latitude | Longitude |
|---|---|---|---|---|
| AHQC_1.113 | progeny phenotyping | A | 44.433051 | -110.81166 |
| AHQC_1.203 | ddRAD genotyping | A | 44.43284 | -110.81212 |
| AHQC_1.209 | ddRAD genotyping | A | 44.432904 | -110.81188 |
| AHQC_1.211 | ddRAD genotyping | A | 44.432916 | -110.81202 |
| AHQC_1.228 | ddRAD genotyping | A | 44.432969 | -110.81188 |
| AHQC_1.234 | ddRAD genotyping | A | 44.432967 | -110.81172 |
| AHQC_1.236 | ddRAD genotyping | A | 44.432985 | -110.81183 |
| AHQC_1.249 | ddRAD genotyping | A | 44.433019 | -110.8118 |
| AHQC_1.251 | ddRAD genotyping | A | 44.43307 | -110.8119 |
| AHQC_1.277 | ddRAD genotyping | A | 44.433093 | -110.81185 |
| AHQC_1.281 | ddRAD genotyping | A | 44.433048 | -110.81161 |
| AHQC_1.29 | progeny phenotyping | A | 44.432904 | -110.81188 |
| AHQC_1.290 | ddRAD genotyping | A | 44.433082 | -110.81172 |
| AHQC_1.297 | ddRAD genotyping | A | 44.433062 | -110.81156 |
| AHQC_1.300 | ddRAD genotyping | A | 44.43309 | -110.81159 |
| AHQC_1.42 | progeny phenotyping | A | 44.432848 | -110.81196 |
| AHQC_1.68 | progeny phenotyping | A | 44.432966 | -110.81204 |
| AHQC_1.92 | progeny phenotyping | A | 44.432918 | -110.81218 |
| AHQNT_1.1 | ddRAD genotyping | P | 44.435226 | -110.80921 |
| AHQNT_1.127 | progeny phenotyping | P | 44.435183 | -110.80917 |
| AHQNT_1.134 | progeny phenotyping | P | 44.43522 | -110.80886 |
| AHQNT_1.136 | progeny phenotyping | P | 44.435116 | -110.80891 |
| AHQNT_1.143 | progeny phenotyping | P | 44.435436 | -110.80835 |
| AHQNT_1.2 | ddRAD genotyping | P | 44.435183 | -110.80917 |
| AHQNT_2.1 | ddRAD genotyping | P | 44.435299 | -110.80895 |
| AHQNT_2.2 | ddRAD genotyping | P | 44.435264 | -110.8089 |
| AHQNT_2.3 | ddRAD genotyping | P | 44.43522 | -110.80886 |
| AHQNT_4.1 | ddRAD genotyping | P | 44.435191 | -110.80839 |
| AHQNT_4.3 | ddRAD genotyping | P | 44.435436 | -110.80835 |
| AHQT_1.13 | progeny phenotyping | A | 44.431354 | -110.81474 |
| AHQT_1.17 | progeny phenotyping | A | 44.432127 | -110.81335 |
| RCG_1.195 | progeny phenotyping | A | 44.519751 | -110.81379 |
| RCG_1.218 | progeny phenotyping | A | 44.518843 | -110.81516 |
| RCG_1.230 | progeny phenotyping | A | 44.518411 | -110.81574 |
| RCG_1.253 | progeny phenotyping | A | 44.518133 | -110.81643 |
| RCG_1.285 | progeny phenotyping | A | 44.517675 | -110.81732 |
| RCG_1.327 | progeny phenotyping | P | 44.523133 | -110.81394 |
| RCG_1.441 | ddRAD genotyping | A | 44.520229 | -110.81315 |
| RCG_1.443 | ddRAD genotyping | A | 44.520045 | -110.81322 |
| RCG_1.451 | ddRAD genotyping | A | 44.519873 | -110.81347 |
| RCG_1.453 | ddRAD genotyping | A | 44.519837 | -110.81367 |
| RCG_1.473 | ddRAD genotyping | A | 44.519213 | -110.81415 |
| RCG_1.477 | ddRAD genotyping | A | 44.519104 | -110.81465 |
| RCG_1.497 | ddRAD genotyping | A | 44.518865 | -110.81502 |

| | | | | |
|---|---|---|---|---|
| RCG_1.513 | ddRAD genotyping | A | 44.518621 | -110.81568 |
| RCG_1.524 | ddRAD genotyping | A | 44.518359 | -110.81587 |
| RCG_1.564 | ddRAD genotyping | A | 44.518133 | -110.81643 |
| RCG_1.570 | ddRAD genotyping | A | 44.518089 | -110.81653 |
| RCG_1.616 | ddRAD genotyping | A | 44.517735 | -110.81717 |
| RCG_1.624 | ddRAD genotyping | A | 44.517675 | -110.81732 |
| RCUV_1.405 | ddRAD genotyping | A | 44.521953 | -110.81079 |
| RCUV_1.409 | ddRAD genotyping | A | 44.521759 | -110.81073 |
| RCUV_1.411 | ddRAD genotyping | A | 44.521804 | -110.81081 |
| RCUV_1.413 | ddRAD genotyping | A | 44.521721 | -110.81085 |
| RCUV_1.421 | ddRAD genotyping | A | 44.521681 | -110.81099 |
| RCUV_1.423 | ddRAD genotyping | A | 44.521625 | -110.81099 |
| RCUV_1.435 | ddRAD genotyping | A | 44.521501 | -110.81112 |
| RCUV_1.438 | ddRAD genotyping | A | 44.521584 | -110.81126 |
| RCUV_1.440 | ddRAD genotyping | A | 44.521481 | -110.81117 |
| YCGP_1.1 | ddRAD genotyping, progeny phenotyping | P | 44.4518 | -110.68537 |
| YGBR_1.2 | progeny phenotyping | P | 44.68405 | -110.74369 |
| YGBR_10.1 | progeny phenotyping | P | 44.70995 | -110.74122 |
| YGBR_10.3 | ddRAD genotyping | P | 44.70995 | -110.74122 |
| YGBR_11.3 | ddRAD genotyping, progeny phenotyping | P | 44.6508 | -110.80226 |
| YGBR_12.1 | ddRAD genotyping, progeny phenotyping | P | 44.65036 | -110.84577 |
| YGBR_13.2 | ddRAD genotyping, progeny phenotyping | P | 44.65018 | -110.84634 |
| YGBR_2.1 | progeny phenotyping | P | 44.68085 | -110.74455 |
| YGBR_2.2 | ddRAD genotyping | P | 44.68085 | -110.74455 |
| YGBR_3.1 | progeny phenotyping | P | 44.68078 | -110.74446 |
| YGBR_3.2 | ddRAD genotyping | P | 44.68078 | -110.74446 |
| YGBR_4.2 | ddRAD genotyping | P | 44.68063 | -110.74457 |
| YGBR_4.3 | progeny phenotyping | P | 44.68063 | -110.74457 |
| YGBR_5.1 | progeny phenotyping | P | 44.68014 | -110.74499 |
| YGBR_6.1 | ddRAD genotyping, progeny phenotyping | A | 44.6802 | -110.74464 |
| YGBR_6.2 | ddRAD genotyping | A | 44.6802 | -110.74464 |
| YGBR_7.1 | ddRAD genotyping, progeny phenotyping | P | 44.68967 | -110.74558 |
| YGBR_7.3 | ddRAD genotyping | P | 44.68967 | -110.74558 |
| YGBR_8.3 | ddRAD genotyping, progeny phenotyping | P | 44.69443 | -110.75999 |
| YGBR_9.1 | ddRAD genotyping, progeny phenotyping | P | 44.69169 | -110.74972 |
| YGBR_9.2 | ddRAD genotyping | P | 44.69169 | -110.74972 |
| YHLB_1.4 | ddRAD genotyping, progeny phenotyping | P | 44.28167 | -110.5064 |
| YHLB_1.5 | ddRAD genotyping | A | 44.28167 | -110.5064 |
| YHLB_10.2 | progeny phenotyping | P | 44.29819 | -110.51722 |
| YHLB_10.4 | ddRAD genotyping | P | 44.29819 | -110.51722 |
| YHLB_11.1 | progeny phenotyping | P | 44.29819 | -110.51722 |
| YHLB_12.3 | progeny phenotyping | P | 44.29901 | -110.51871 |
| YHLB_13.3 | ddRAD genotyping, progeny phenotyping | P | 44.30096 | -110.51966 |
| YHLB_14.1 | ddRAD genotyping | A | 44.30876 | -110.52599 |
| YHLB_14.5 | progeny phenotyping | A | 44.30876 | -110.52599 |

| | | | | |
|---|---|---|---|---|
| YHLB_15.1 | progeny phenotyping | A | 44.3134 | -110.53607 |
| YHLB_2.1 | ddRAD genotyping, progeny phenotyping | P | 44.28149 | -110.50646 |
| YHLB_2.2 | ddRAD genotyping | P | 44.28149 | -110.50646 |
| YHLB_3.3 | ddRAD genotyping | A | 44.28164 | -110.507 |
| YHLB_3.4 | ddRAD genotyping, progeny phenotyping | A | 44.28164 | -110.507 |
| YHLB_4.1 | ddRAD genotyping | P | 44.28168 | -110.507 |
| YHLB_4.2 | progeny phenotyping | P | 44.28168 | -110.507 |
| YHLB_5.2 | progeny phenotyping | P | 44.28288 | -110.50556 |
| YHLB_6.1 | progeny phenotyping | P | 44.28996 | -110.50391 |
| YHLB_6.3 | ddRAD genotyping | P | 44.28996 | -110.50391 |
| YHLB_7.2 | ddRAD genotyping, progeny phenotyping | A | 44.29128 | -110.50526 |
| YHLB_8.7 | ddRAD genotyping | A | 44.29087 | -110.5065 |
| YHLB_9.2 | ddRAD genotyping, progeny phenotyping | A | 44.29069 | -110.50654 |
| YLGB_1.1 | ddRAD genotyping, progeny phenotyping | A | 44.53564 | -110.79978 |
| YLGB_1.2 | ddRAD genotyping | A | 44.53564 | -110.79978 |
| YLGB_10.3 | progeny phenotyping | A | 44.54433 | -110.78584 |
| YLGB_11.4 | progeny phenotyping | A | 44.54451 | -110.78661 |
| YLGB_13.2 | progeny phenotyping | P | 44.54418 | -110.78792 |
| YLGB_14.3 | ddRAD genotyping, progeny phenotyping | A | 44.54485 | -110.78782 |
| YLGB_15.1 | progeny phenotyping | A | 44.54387 | -110.78472 |
| YLGB_16.2 | progeny phenotyping | A | 44.54411 | -110.78413 |
| YLGB_17.3 | ddRAD genotyping | A | 44.54237 | -110.7907 |
| YLGB_18.1 | ddRAD genotyping | P | 44.55012 | -110.80646 |
| YLGB_18.2 | progeny phenotyping | P | 44.55012 | -110.80646 |
| YLGB_19.1 | ddRAD genotyping | A | 44.5511 | -110.80717 |
| YLGB_19.3 | progeny phenotyping | A | 44.5511 | -110.80717 |
| YLGB_2.2 | ddRAD genotyping, progeny phenotyping | A | 44.53483 | -110.8002 |
| YLGB_2.3 | ddRAD genotyping | A | 44.53483 | -110.8002 |
| YLGB_20.2 | progeny phenotyping | A | 44.55013 | -110.80604 |
| YLGB_21.2 | progeny phenotyping | A | 44.55015 | -110.80827 |
| YLGB_22.1 | progeny phenotyping | A | 44.5501 | -110.80827 |
| YLGB_23.2 | ddRAD genotyping | P | 44.53544 | -110.80828 |
| YLGB_23.3 | ddRAD genotyping, progeny phenotyping | P | 44.53544 | -110.80828 |
| YLGB_24.1 | ddRAD genotyping | P | 44.5335 | -110.80718 |
| YLGB_24.3 | ddRAD genotyping, progeny phenotyping | P | 44.5335 | -110.80718 |
| YLGB_25.1 | progeny phenotyping | A | 44.53369 | -110.80693 |
| YLGB_27.1 | ddRAD genotyping, progeny phenotyping | P | 44.53376 | -110.80571 |
| YLGB_27.3 | ddRAD genotyping | P | 44.53376 | -110.80571 |
| YLGB_28.1 | ddRAD genotyping | P | 44.53387 | -110.80557 |
| YLGB_28.3 | ddRAD genotyping, progeny phenotyping | P | 44.53387 | -110.80557 |
| YLGB_29.2 | ddRAD genotyping | P | 44.53446 | -110.80754 |
| YLGB_29.3 | progeny phenotyping | P | 44.53446 | -110.80754 |
| YLGB_3.1 | ddRAD genotyping | A | 44.53484 | -110.8003 |
| YLGB_3.2 | progeny phenotyping | A | 44.53484 | -110.8003 |
| YLGB_30.4 | ddRAD genotyping | P | 44.53413 | -110.79808 |

| | | | | |
|---|---|---|---|---|
| YLGB_31.1 | progeny phenotyping | P | 44.53288 | -110.79717 |
| YLGB_31.2 | ddRAD genotyping | P | 44.53288 | -110.79717 |
| YLGB_4.2 | progeny phenotyping | P | 44.53504 | -110.80087 |
| YLGB_5.1 | progeny phenotyping | A | 44.539 | -110.80242 |
| YLGB_6.1 | ddRAD genotyping | A | 44.53919 | -110.80304 |
| YLGB_7.3 | progeny phenotyping | A | 44.53922 | -110.8041 |
| YLGB_8.2 | progeny phenotyping | A | 44.53958 | -110.80161 |
| YLGB_9.1 | progeny phenotyping | A | 44.53911 | -110.80161 |
| YLSG_1.3 | progeny phenotyping | A | 44.41676 | -110.80863 |
| YLSG_2.1 | progeny phenotyping | A | 44.41638 | -110.80908 |
| YLSG_3.1 | ddRAD genotyping, progeny phenotyping | P | 44.41816 | -110.80629 |
| YLSG_3.2 | ddRAD genotyping | P | 44.41816 | -110.80629 |
| YLSG_4.4 | ddRAD genotyping, progeny phenotyping | A | 44.4167 | -110.81339 |
| YLSG_5.4 | ddRAD genotyping, progeny phenotyping | P | 44.4164 | -110.81376 |
| YLSG_6.2 | ddRAD genotyping | P | 44.41512 | -110.8127 |
| YLSG_6.3 | progeny phenotyping | P | 44.41512 | -110.8127 |
| YLSG_7.3 | progeny phenotyping | A | 44.41561 | -110.81162 |
| YLSG_8.1 | progeny phenotyping | A | 44.41587 | -110.80976 |
| YLSG_9.3 | progeny phenotyping | A | 44.41575 | -110.80962 |
| YMAR_1.2 | ddRAD genotyping, progeny phenotyping | P | 44.64207 | -110.87631 |
| YMAR_3.1 | ddRAD genotyping | P | 44.64366 | -110.91594 |
| YMAR_3.2 | ddRAD genotyping, progeny phenotyping | P | 44.64366 | -110.91594 |
| YMGB_1.1 | ddRAD genotyping, progeny phenotyping | A | 44.5152 | -110.8323 |
| YMGB_1.2 | ddRAD genotyping | A | 44.5152 | -110.8323 |
| YMGB_10.1 | ddRAD genotyping | A | 44.5271 | -110.83662 |
| YMGB_10.3 | ddRAD genotyping, progeny phenotyping | A | 44.5271 | -110.83662 |
| YMGB_11.1 | ddRAD genotyping, progeny phenotyping | A | 44.52715 | -110.83678 |
| YMGB_11.3 | ddRAD genotyping | A | 44.52715 | -110.83678 |
| YMGB_12.3 | progeny phenotyping | A | 44.5267 | -110.83693 |
| YMGB_13.3 | progeny phenotyping | A | 44.52787 | -110.83662 |
| YMGB_13.9 | ddRAD genotyping | A | 44.52787 | -110.83662 |
| YMGB_2.1 | ddRAD genotyping | A | 44.51608 | -110.83267 |
| YMGB_2.3 | ddRAD genotyping, progeny phenotyping | A | 44.51608 | -110.83267 |
| YMGB_3.2 | ddRAD genotyping, progeny phenotyping | A | 44.51715 | -110.83205 |
| YMGB_3.4 | ddRAD genotyping | A | 44.51715 | -110.83205 |
| YMGB_5.2 | ddRAD genotyping, progeny phenotyping | A | 44.51793 | -110.83089 |
| YMGB_6.1 | progeny phenotyping | A | 44.51635 | -110.83296 |
| YMGB_6.3 | ddRAD genotyping | A | 44.51635 | -110.83296 |
| YMGB_7.1 | progeny phenotyping | P | 44.51613 | -110.83301 |
| YMGB_8.2 | ddRAD genotyping | A | 44.51413 | -110.83442 |
| YMGB_8.3 | progeny phenotyping | A | 44.51413 | -110.83442 |
| YMGB_9.1 | progeny phenotyping | A | 44.51459 | -110.83415 |
| YMGB_9.2 | ddRAD genotyping | A | 44.51459 | -110.83415 |
| YNGB_1.1 | ddRAD genotyping, progeny phenotyping | P | 44.71668 | -110.71449 |
| YNGB_1.3 | ddRAD genotyping | P | 44.71668 | -110.71449 |

| | | | | |
|---|---|---|---|---|
| YOJO_1.5 | progeny phenotyping | A | 44.5629 | -110.8387 |
| YOJO_2.2 | progeny phenotyping | A | 44.56269 | -110.8389 |
| YOJO_3.2 | progeny phenotyping | A | 44.56292 | -110.83895 |
| YOJO_4.1 | progeny phenotyping | A | 44.56162 | -110.83582 |
| YOJO_5.1 | progeny phenotyping | A | 44.5613 | -110.83534 |
| YOJO_6.3 | progeny phenotyping | P | 44.56116 | -110.83489 |
| YOJO_7.1 | progeny phenotyping | P | 44.56063 | -110.83457 |
| YOJO_7.3 | ddRAD genotyping | P | 44.56063 | -110.83457 |
| YOJO_8.3 | ddRAD genotyping, progeny phenotyping | A | 44.65073 | -110.83466 |
| YSGB_1.4 | progeny phenotyping | A | 44.35645 | -110.79768 |
| YSGB_10.1 | ddRAD genotyping | P | 44.35146 | -110.79825 |
| YSGB_10.2 | ddRAD genotyping, progeny phenotyping | P | 44.35146 | -110.79825 |
| YSGB_11.1 | ddRAD genotyping, progeny phenotyping | P | 44.35341 | -110.79863 |
| YSGB_12.1 | ddRAD genotyping | P | 44.36125 | -110.79777 |
| YSGB_12.2 | ddRAD genotyping | P | 44.36125 | -110.79777 |
| YSGB_13.1 | ddRAD genotyping, progeny phenotyping | P | 44.3637 | -110.79762 |
| YSGB_13.2 | ddRAD genotyping | P | 44.3637 | -110.79762 |
| YSGB_15.2 | ddRAD genotyping | P | 44.36633 | -110.80172 |
| YSGB_15.3 | ddRAD genotyping | P | 44.36633 | -110.80172 |
| YSGB_16.3 | ddRAD genotyping | P | 44.36577 | -110.80597 |
| YSGB_17.4 | ddRAD genotyping | P | 44.36574 | -110.80612 |
| YSGB_18.2 | ddRAD genotyping | P | 44.3694 | -110.8122 |
| YSGB_18.3 | ddRAD genotyping | P | 44.3694 | -110.8122 |
| YSGB_19.1 | progeny phenotyping | P | 44.37069 | -110.81286 |
| YSGB_2.2 | ddRAD genotyping, progeny phenotyping | A | 44.35524 | -110.79784 |
| YSGB_2.4 | ddRAD genotyping | A | 44.35524 | -110.79784 |
| YSGB_20.2 | progeny phenotyping | P | 44.37086 | -110.81293 |
| YSGB_20.4 | ddRAD genotyping | P | 44.37086 | -110.81293 |
| YSGB_21.1 | ddRAD genotyping | P | 44.37166 | -110.81274 |
| YSGB_22.1 | ddRAD genotyping, progeny phenotyping | P | 44.40482 | -110.82713 |
| YSGB_22.2 | ddRAD genotyping | P | 44.40482 | -110.82713 |
| YSGB_23.1 | ddRAD genotyping, progeny phenotyping | P | 44.43478 | -110.80538 |
| YSGB_3.2 | ddRAD genotyping, progeny phenotyping | P | 44.35529 | -110.79784 |
| YSGB_3.3 | ddRAD genotyping | P | 44.35529 | -110.79784 |
| YSGB_4.4 | ddRAD genotyping, progeny phenotyping | A | 44.35484 | -110.79906 |
| YSGB_5.1 | ddRAD genotyping, progeny phenotyping | P | 44.35411 | -110.79845 |
| YSGB_5.3 | ddRAD genotyping | P | 44.35411 | -110.79845 |
| YSGB_6.1 | ddRAD genotyping, progeny phenotyping | P | 44.35378 | -110.79861 |
| YSGB_6.3 | ddRAD genotyping | P | 44.35378 | -110.79861 |
| YSGB_7.2 | ddRAD genotyping, progeny phenotyping | A | 44.35243 | -110.79788 |
| YSGB_7.4 | ddRAD genotyping | A | 44.35243 | -110.79788 |
| YSGB_8.1 | ddRAD genotyping | P | 44.35206 | -110.8007 |
| YSGB_8.2 | ddRAD genotyping, progeny phenotyping | P | 44.35206 | -110.8007 |
| YSGB_9.4 | ddRAD genotyping, progeny phenotyping | A | 44.35172 | -110.80007 |
| YSGB_9.5 | ddRAD genotyping | A | 44.35172 | -110.80007 |

| | | | | |
|---|---|---|---|---|
| YUGB_1.1 | ddRAD genotyping | P | 44.46266 | -110.82784 |
| YUGB_1.2 | ddRAD genotyping, progeny phenotyping | P | 44.46266 | -110.82784 |
| YUGB_10.1 | progeny phenotyping | A | 44.46513 | -110.83701 |
| YUGB_10.2 | ddRAD genotyping | A | 44.46513 | -110.83701 |
| YUGB_11.2 | ddRAD genotyping | A | 44.46594 | -110.83645 |
| YUGB_12.1 | ddRAD genotyping | P | 44.46899 | -110.8406 |
| YUGB_12.3 | ddRAD genotyping | P | 44.46899 | -110.8406 |
| YUGB_13.1 | ddRAD genotyping | A | 44.46898 | -110.84055 |
| YUGB_13.2 | ddRAD genotyping, progeny phenotyping | A | 44.46898 | -110.84055 |
| YUGB_14.1 | progeny phenotyping | A | 44.47162 | -110.84155 |
| YUGB_15.1 | progeny phenotyping | A | 44.47285 | -110.84151 |
| YUGB_15.2 | ddRAD genotyping | A | 44.47285 | -110.84151 |
| YUGB_16.1 | ddRAD genotyping, progeny phenotyping | P | 44.47364 | -110.84141 |
| YUGB_16.2 | ddRAD genotyping | P | 44.47364 | -110.84141 |
| YUGB_17.1 | ddRAD genotyping | P | 44.47446 | -110.84279 |
| YUGB_17.2 | progeny phenotyping | P | 44.47446 | -110.84279 |
| YUGB_18.1 | ddRAD genotyping | P | 44.47451 | -110.8432 |
| YUGB_18.2 | ddRAD genotyping, progeny phenotyping | P | 44.47451 | -110.8432 |
| YUGB_19.1 | ddRAD genotyping | P | 44.47575 | -110.84433 |
| YUGB_19.2 | ddRAD genotyping, progeny phenotyping | P | 44.47575 | -110.84433 |
| YUGB_2.1 | ddRAD genotyping | A | 44.46374 | -110.82833 |
| YUGB_2.2 | ddRAD genotyping, progeny phenotyping | A | 44.46374 | -110.82833 |
| YUGB_20.1 | ddRAD genotyping, progeny phenotyping | A | 44.47925 | -110.8489 |
| YUGB_20.3 | ddRAD genotyping | A | 44.47925 | -110.8489 |
| YUGB_21.2 | ddRAD genotyping, progeny phenotyping | A | 44.47975 | -110.85053 |
| YUGB_22.1 | ddRAD genotyping, progeny phenotyping | P | 44.48028 | -110.84914 |
| YUGB_22.2 | ddRAD genotyping | P | 44.48028 | -110.84914 |
| YUGB_23.1 | ddRAD genotyping | P | 44.47966 | -110.84891 |
| YUGB_23.2 | ddRAD genotyping, progeny phenotyping | P | 44.47966 | -110.84891 |
| YUGB_24.1 | ddRAD genotyping, progeny phenotyping | P | 44.46165 | -110.8543 |
| YUGB_24.2 | ddRAD genotyping | P | 44.46165 | -110.8543 |
| YUGB_25.2 | ddRAD genotyping, progeny phenotyping | A | 44.46138 | -110.85496 |
| YUGB_26.1 | ddRAD genotyping, progeny phenotyping | P | 44.46097 | -110.85568 |
| YUGB_26.2 | ddRAD genotyping | P | 44.46097 | -110.85568 |
| YUGB_27.1 | ddRAD genotyping, progeny phenotyping | A | 44.46267 | -110.85499 |
| YUGB_27.2 | ddRAD genotyping | A | 44.46267 | -110.85499 |
| YUGB_28.3 | ddRAD genotyping, progeny phenotyping | A | 44.48562 | -110.85368 |
| YUGB_29.3 | ddRAD genotyping | A | 44.48577 | -110.85315 |
| YUGB_29.4 | ddRAD genotyping, progeny phenotyping | A | 44.48577 | -110.85315 |
| YUGB_3.1 | ddRAD genotyping | A | 44.46477 | -110.82979 |
| YUGB_3.2 | ddRAD genotyping, progeny phenotyping | A | 44.46477 | -110.82979 |
| YUGB_30.1 | ddRAD genotyping | A | 44.48582 | -110.85315 |
| YUGB_30.2 | ddRAD genotyping | A | 44.48582 | -110.85315 |
| YUGB_30.3 | progeny phenotyping | A | 44.48582 | -110.85315 |
| YUGB_31.2 | progeny phenotyping | A | 44.48512 | -110.85478 |

| | | | | |
|---|---|---|---|---|
| YUGB_32.5 | ddRAD genotyping | A | 44.48591 | -110.85609 |
| YUGB_33.3 | ddRAD genotyping, progeny phenotyping | A | 44.48648 | -110.85645 |
| YUGB_33.5 | ddRAD genotyping | A | 44.48648 | -110.85645 |
| YUGB_34.3 | progeny phenotyping | A | 44.48573 | -110.85751 |
| YUGB_35.1 | progeny phenotyping | A | 44.48463 | -110.85611 |
| YUGB_4.1 | ddRAD genotyping | P | 44.46305 | -110.8305 |
| YUGB_4.2 | ddRAD genotyping, progeny phenotyping | P | 44.46305 | -110.8305 |
| YUGB_5.1 | ddRAD genotyping | A | 44.4629 | -110.82878 |
| YUGB_5.1 | progeny phenotyping | A | 44.4629 | -110.82878 |
| YUGB_5.2 | ddRAD genotyping | A | 44.4629 | -110.82878 |
| YUGB_6.1 | progeny phenotyping | P | 44.46193 | -110.82785 |
| YUGB_7.1 | ddRAD genotyping | A | 44.46193 | -110.82903 |
| YUGB_7.2 | ddRAD genotyping, progeny phenotyping | A | 44.46193 | -110.82903 |
| YUGB_8.1 | ddRAD genotyping | A | 44.46486 | -110.83705 |
| YUGB_8.2 | ddRAD genotyping, progeny phenotyping | A | 44.46486 | -110.83705 |
| YUGB_9.1 | ddRAD genotyping, progeny phenotyping | A | 44.46483 | -110.83705 |
| YUGB_9.3 | ddRAD genotyping | A | 44.46483 | -110.83705 |
| YVTC_1.1 | ddRAD genotyping | P | 44.6533 | -110.5456 |
| YVTC_1.2 | ddRAD genotyping, progeny phenotyping | P | 44.6533 | -110.5456 |
| YVTC_10.2 | ddRAD genotyping, progeny phenotyping | P | 44.64907 | -110.55041 |
| YVTC_10.3 | ddRAD genotyping | P | 44.64907 | -110.55041 |
| YVTC_11.1 | ddRAD genotyping | P | 44.65504 | -110.54228 |
| YVTC_11.3 | progeny phenotyping | P | 44.65504 | -110.54228 |
| YVTC_2.1 | ddRAD genotyping | P | 44.65226 | -110.5452 |
| YVTC_2.2 | ddRAD genotyping, progeny phenotyping | P | 44.65226 | -110.5452 |
| YVTC_3.2 | progeny phenotyping | A | 44.65064 | -110.55774 |
| YVTC_4.1 | progeny phenotyping | A | 44.65118 | -110.55791 |
| YVTC_5.1 | ddRAD genotyping | P | 44.65123 | -110.55795 |
| YVTC_5.3 | ddRAD genotyping, progeny phenotyping | P | 44.65123 | -110.55795 |
| YVTC_6.1 | ddRAD genotyping, progeny phenotyping | A | 44.65031 | -110.55688 |
| YVTC_6.3 | ddRAD genotyping | A | 44.65031 | -110.55688 |
| YVTC_7.1 | ddRAD genotyping | A | 44.65005 | -110.55637 |
| YVTC_7.2 | ddRAD genotyping, progeny phenotyping | A | 44.65005 | -110.55637 |
| YVTC_8.2 | ddRAD genotyping, progeny phenotyping | P | 44.65033 | -110.55583 |
| YVTC_9.3 | ddRAD genotyping, progeny phenotyping | P | 44.65058 | -110.55493 |
| YWTB_1.1 | ddRAD genotyping | A | 44.4157 | -110.57109 |
| YWTB_1.2 | ddRAD genotyping, progeny phenotyping | A | 44.4157 | -110.57109 |
| YWTB_2.1 | ddRAD genotyping | A | 44.41591 | -110.56966 |
| YWTB_2.2 | ddRAD genotyping, progeny phenotyping | A | 44.41591 | -110.56966 |
| YWTB_3.2 | ddRAD genotyping | P | 44.41591 | -110.56966 |
| YWTB_3.3 | progeny phenotyping | P | 44.41591 | -110.56966 |
| YWTB_4.1 | ddRAD genotyping | A | 44.41825 | -110.57205 |
| YWTB_4.2 | ddRAD genotyping, progeny phenotyping | A | 44.41825 | -110.57205 |
| YWTB_5.2 | ddRAD genotyping, progeny phenotyping | A | 44.41899 | -110.57193 |
| YWTB_6.2 | progeny phenotyping | P | 44.41709 | -110.57317 |

| | | | | |
|---|---|---|---|---|
| YWTB_6.4 | ddRAD genotyping | P | 44.41709 | -110.57317 |
| YWTB_7.1 | ddRAD genotyping | P | 44.41642 | -110.57293 |
| YWTB_8.2 | progeny phenotyping | A | 44.41563 | -110.57236 |
| YWTB_9.3 | progeny phenotyping | A | 44.41669 | -110.57187 |
